# Where is all the nonlinearity: flexible nonlinear modeling of behaviorally relevant neural dynamics using recurrent neural networks

**DOI:** 10.1101/2021.09.03.458628

**Authors:** Omid G. Sani, Bijan Pesaran, Maryam M. Shanechi

## Abstract

Understanding the dynamical transformation of neural activity to behavior requires modeling this transformation while both dissecting its potential nonlinearities and dissociating and preserving its nonlinear behaviorally relevant neural dynamics, which remain unaddressed. We present RNN PSID, a nonlinear dynamic modeling method that enables flexible dissection of nonlinearities, dissociation and preferential learning of neural dynamics relevant to specific behaviors, and causal decoding. We first validate RNN PSID in simulations and then use it to investigate nonlinearities in monkey spiking and LFP activity across four tasks and different brain regions. Nonlinear RNN PSID successfully dissociated and preserved nonlinear behaviorally relevant dynamics, thus outperforming linear and non-preferential nonlinear learning methods in behavior decoding while reaching similar neural prediction. Strikingly, dissecting the nonlinearities with RNN PSID revealed that consistently across all tasks, summarizing the nonlinearity only in the mapping from the latent dynamics to behavior was largely sufficient for predicting behavior and neural activity. RNN PSID provides a novel tool to reveal new characteristics of nonlinear neural dynamics underlying behavior.

## Introduction

Understanding the dynamics of neural population activity and how they give rise to behavior is a major goal across diverse domains of neuroscience and neuroengineering. Toward this goal, dynamic models of neural population activity describe it in terms of a low-dimensional latent state embedded in the high-dimensional space of neural recordings, while also describing the temporal structure of the state evolution in its low-dimensional subspace and subsequently relating the latent state to behavior^1–6^. This approach ultimately models a dynamical transformation from neural activity to behavior. However, precise modeling remains quite challenging because the dynamical transformation of neural activity to behavior can exhibit nonlinearities, these nonlinearities may be introduced in one or more different elements within this transformation—e.g., in the evolution of the state or in its embedding—, and finally these dynamics may relate to a multitude of behaviors and/or internal states simultaneously^7,8^. To date, developing a unified dynamic modeling framework that can capture nonlinearities in behaviorally relevant neural dynamics, dissect and discover the origin of these nonlinearities, and finally preferentially learn and dissociate these nonlinear behaviorally relevant dynamics from other neural dynamics has remained elusive.

Prior dynamic models of neural population dynamics have often been linear or generalized linear^1,4,9–12^. Thus, recently there has been growing interest toward models that support piece-wise linear^13^, switching linear^5,14^, or nonlinear^2,15–18^ neural dynamics, especially in applications such as single trial smoothing of neural population activity^2^ and decoding behavior^15,16,18^. However, two major challenges have remained unaddressed in modeling the dynamical transformation of neural activity to behavior. First, these nonlinear models do not dissect the origin of nonlinearities in this transformation. Indeed, this transformation can be decomposed into several interpretable elements (**Fig. 1a,b**): the mapping from neural activity to the latent subspace (neural drive), the state dynamics in this subspace (recursion), and finally the mappings of the state to neural activity and behavior (neural and behavior readouts). Dissecting the origin of nonlinearity requires novel dynamic modeling methods that can explicitly represent each of these interpretable elements and flexibly make each individual element linear or nonlinear. Second, given that neural activity can simultaneously be related to multiple behaviors and/or internal states^7,8^, understanding this transformation requires learning models of nonlinear neural dynamics while also prioritizing the learning of those dynamics that are related to the specific behavior of interest and dissociating them from other neural dynamics^1^. To date, this prioritized dissociation has only been possible for linear dynamic models through a method termed PSID that uses linear algebraic subspace identification principles^1^, but remains unaddressed for nonlinear models. Here, we address both challenges within a novel unified nonlinear dynamic modeling framework using recurrent neural networks (RNNs), while also enabling causal and efficient nonlinear decoding of the latent states and behavior from neural dynamics.

**Fig. 1.**
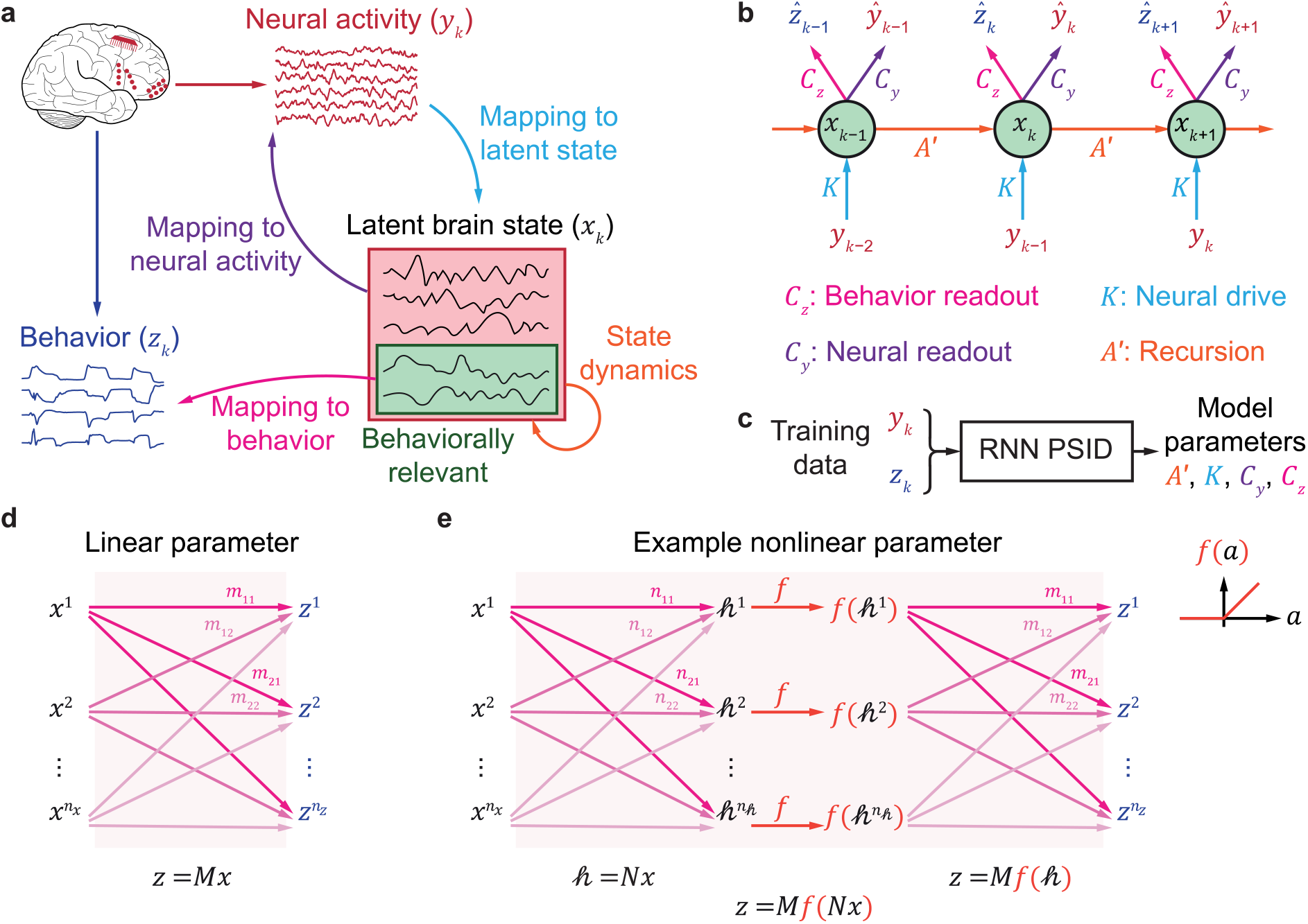
High-level view of RNN PSID. (**a**) RNN PSID aims to learn the mapping of neural activity (*y*_*k*_) to latent states (*x*_*k*_), learn the temporal structure of the latent states (i.e. state dynamics), dissociate the behaviorally relevant latent states that are relevant to any measured behavior (*z*_*k*_) from other states, and learn the mapping of the latent states to behavior and to neural activity, while allowing flexible linear or nonlinear mappings in any of these elements. RNN PSID additionally aims to prioritize the learning of behaviorally relevant neural dynamics while also optionally allowing the learning of other neural dynamics. (**b**) Computation graph of the RNN PSID model consists of an RNN with neural activity at the current time step as the input and with the behavior and neural activity in the next time step as the outputs (Methods). Each mapping element from (a) has a corresponding parameter in the model indicated by the same colors and termed neural drive, recursion, behavior readout, and neural readout. For the complete two-stage computation graph see **S Fig. 1**. (**c**) PSID learns all model parameters using training neural and behavior data. After a model is learned, only past neural activity is used to decode behavior and predict neural activity using the computation graph in (b). (**d,e**) Any one or all of the model parameters from (b) can be either a linear matrix (panel d), or in general an arbitrary multilayer feed-forward neural network (panel e). The example feed-forward network in (e) has one hidden layer with *h* units and uses a rectified linear unit (ReLU) activation function for the hidden layer.

RNNs provide an especially potent architecture for building nonlinear dynamic models^2,15,16,19^. A major part of the appeal of RNNs is that regardless of the computational graph of the model, model parameters can always be learned from the training data using general numerical optimization approaches^20^, which can enable easy exploration of different model structures. Several prior works have used RNNs to build nonlinear dynamic models, either causally^15,16^ or non-causally^2,3^. However, none of these works aim to enable dissociation of behaviorally relevant neural dynamics from other neural dynamics or dissect nonlinearities to probe characteristics such as the origin of nonlinearities, leaving both of the above challenges unaddressed.

Here, we develop a new nonlinear dynamic modeling method termed RNN preferential system identification (PSID), or RNN PSID, that addresses both challenges while enabling causal decoding. Three key ideas enable these capabilities in RNN PSID. First, we use RNNs trained using numerical optimization, which results in a flexible nonlinear dynamic model whose components correspond to different interpretable elements within the dynamical transformation of neural activity to behavior, and can each be flexibly chosen to have various nonlinear or linear structures (**Fig. 1b-e**). This flexibility enables dissecting the type and origin of nonlinearity and thus addresses the first challenge. Second, we devise a two-stage learning approach^1^ where two stacked RNNs are learned with appropriate distinct optimization objectives, such that one RNN learns behaviorally relevant neural dynamics with priority, and then the other RNN learns any remaining neural dynamics (**Fig. 1a** and **S Fig. 1**). This allows dissociation of behaviorally relevant neural dynamics from other dynamics while simultaneously enabling the learning of all neural dynamics, thus addressing the second challenge. Third, we formulate the problem in predictor form^21^ such that the inference model is directly learned as the RNNs, thus enabling causal and computationally efficient nonlinear decoding.

We first validate RNN PSID in numerical simulations with known nonlinearities and demonstrate that it successfully learns the nonlinearity in behaviorally relevant dynamics and correctly determines the origin of nonlinearities. We next use RNN PSID to investigate nonlinearities in the dynamics of population spiking activity and local field potential (LFP) modalities in monkeys during four diverse behavioral tasks with recordings from different brain regions. Across all tasks, brain regions, and neural modalities, nonlinear RNN PSID models explain the behaviorally relevant neural dynamics significantly more accurately than linear PSID models, thus suggesting that nonlinearity exists and can be captured by RNN PSID. Moreover, RNN PSID not only extracts the behaviorally relevant neural dynamics more accurately than nonlinear RNN models without prioritization, but also at least as accurately describes the overall neural dynamics regardless of their relevance to behavior. We next dissect the nonlinearities and find their origin in each dataset by comparing RNN PSID models with nonlinearity in different interpretable model components. Strikingly, despite the diversity of tasks and neural recordings and modalities, models that summarize all nonlinearities in the behavior readout from the latent state best explain all four datasets in terms of predicting behavior and neural activity. Together, these results highlight RNN PSID as a novel tool to model, dissociate and prioritize, and dissect nonlinear behaviorally relevant neural dynamics, and reveal consistent new insights about the nonlinearity in these dynamics across four behavioral tasks.

## Results

### Overview of RNN PSID

As a nonlinear generalization of linear state-space models (Methods, **S Note 1**), we model neural activity and behavior jointly as

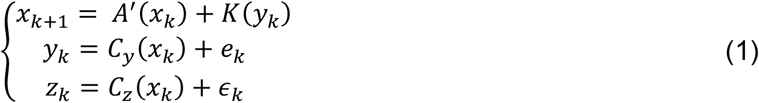

where *k* is the time index, 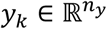 and 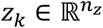 denote the neural activity and behavior time series respectively, 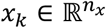 is the latent state, and *e*_*k*_ and *∈*_*k*_ denote neural and behavior dynamics that are unpredictable from past neural activity. Multi-input-multi-output functions *A*′, *K*, *C*_*y*_, and *C*_*z*_ are model parameters that fully specify the model, and have interpretable descriptions (Methods, **Fig. 1a-b**): *K* is the neural drive and specifies the mapping from neural observations to the latent subspace over which the state evolves. *A*′ (recursion) specifies how the latent state moves from one time step to the next, thus specifying its recurrent dynamics. *C*_*y*_ and *C*_*z*_ are the neural and behavior readouts, specifying the mapping from the latent states to neural and behavior observations, respectively. We learn the model parameters using training neural and behavior data via numerical optimization (Methods, **Fig. 1c**). This allows the flexibility of being able to design each of the model parameters (e.g. *C*_*z*_) to be an arbitrary multi-layer neural network (**Fig. 1e**), which as universal approximators can approximate any smooth nonlinear function^22–24^. Overall, equation (1) formulates an RNN (**Fig. 1b**), which under mild conditions can approximate any state-space dynamics^25^. As a special case, if all parameters are set to be linear matrix multiplications (i.e. a fully connected neural network with no-hidden layer, **Fig. 1d**), equation (1) reduces to a standard linear state-space model written in the predictor form^21^ where *x*_*k*_ is the Kalman estimation of the latent states (**S Note 1**). Critically, since the RNN in equation (1) is a nonlinear generalization of this predictor form, once the RNN model parameters are learned from training data, the decoding problem of estimating the latent states from the neural activity is readily solved by iterating through equation (1) and can thus be done causally (Methods).

Our goal is to learn both the nonlinear dynamical transformation of neural population activity to behavior and the nonlinear neural dynamics that are unrelated to the measured behavior. As such, we need a new learning algorithm for the nonlinear state-space model above that enables the dissociation and prioritized learning of behaviorally relevant neural dynamics (**Fig. 1a**) while also being able to describe any other residual neural dynamics. Thus, we devise a two-stage learning procedure consisting of two stacked RNNs (**S Fig. 1**). In the first stage, we learn behaviorally relevant neural dynamics by fitting an RNN model as in equation (1), but by finding the model parameters with the sole objective of numerical optimization being the maximization of the behavior prediction accuracy. The latent state of this RNN, denoted by 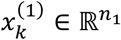, learns the behaviorally relevant neural dynamics. We then learn the mapping between this state and neural activity (denoted by 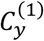) as a feed-forward neural network that optimizes the neural prediction accuracy (Methods). In the second stage, which is optional, we learn any other residual neural dynamics not learned in the first stage by fitting a second RNN, this time with the input being not only the neural activity *y*_*k*_ but also the extracted latent state of the first RNN (i.e. 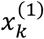). Model parameters of the second RNN are found with the sole objective of numerical optimization being to maximize the prediction accuracy of those neural dynamics that are not predicted by the first RNN (Methods). We then learn the mapping 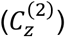 between the latent state of the second RNN denoted by 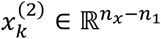 and any behavior dynamics not already predicted by the first RNN as a feed-forward neural network that optimizes the behavior prediction accuracy—this step may be needed if the first RNN state dimension is desired to be very small, or to just perform modeling of all neural dynamics without prioritization using stage 2 alone. Together, the two stages learn two stacked RNNs that predict both behavior and neural activity using past neural activity (**S Fig. 1**). Importantly, either stage could also be used alone depending on the desired goal. If desired, the first stage alone could be used (i.e. *n*_1_ = *n*_*x*_) to only learn behaviorally relevant neural dynamics. Alternatively, the special case in which only the second stage is used (i.e. *n*_1_ = 0) reduces to nonlinear neural dynamic modeling (NDM)^1^ that models neural dynamics irrespective of relevance to behavior, i.e. is non-preferential. We refer to the latter case with only the second stage as RNN NDM.

The flexibility over which model parameters are nonlinear, and further the direct interpretability of these parameters in the process of generating neural and behavior dynamics enables two important capabilities. First, we can search over various combinations of nonlinearities to find the best model according to the training data and use it in the test data (Methods). This will be referred to as flexible nonlinearity and allows us to automatically select the best model for each dataset and get an estimate of the peak performance. Second, we can dissect nonlinearities by comparing alternative models that each set a different individual parameter to be nonlinear or linear independently of other parameters. This analysis finds the single parameter that best summarizes most nonlinearities in the data, and can use the interpretation of this parameter to test hypotheses about where nonlinearities lie in the process of generating neural and behavior dynamics. As a baseline to interpret the nonlinear results, we can set all parameters to be linear, which we refer to as linear RNN PSID or linear PSID for short.

### Model assessment and dissection

To quantify how accurately a learned model describes the behaviorally relevant neural dynamics, we use the model to causally decode behavior using past neural activity. Similarly, to quantify how accurately the model describes the neural activity in general, we use the model to causally predict the neural activity at each time point using past neural activity, which will be referred to as neural self-prediction. Both performance measures are always computed with cross-validation (Methods).

When comparing models, for example to find the best individual nonlinearity to determine the origin of nonlinearity, those models that reach both better behavior decoding and better neural self-prediction will be considered as more accurate representations of data. There may be trade-off cases where one model has better behavior decoding and the other has better neural self-prediction. In such cases, there may not be any clear best model for the data overall, and either model may be more useful depending on the application. To interpret such cases, we use the term “performance frontier” to refer to the range of performances achievable by all the models that are in some sense better than or at least equivalent to every other model. More precisely, when comparing a group 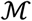 of models, model 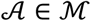 will be described as reaching the performance frontier when compared with every other model 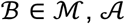 is significantly better than 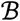 in decoding, or self-prediction, or is comparable to 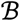 in both. Note that 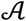 may be better than some model 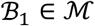 in decoding while being better than another model 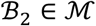 in self-prediction, nevertheless it will be on the frontier as long as in every comparison, there is at least one measure for which 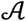 is being more performant or it is at least equally performant in both measures.

### Numerical simulations validate RNN PSID

We first validated RNN PSID with numerical simulations. To confirm the validity of the two-stage numerical optimization approach, we started by focusing on the case where all RNN parameters were linear functions. We generated random realizations from 100 random linear models and used linear RNN PSID to learn the models given the training data time series. In each random model, only a subset of state dimensions contributed to generating behavior and thus were behaviorally relevant (Methods). We computed the behavior decoding and neural self-prediction performance measures for the true models to quantify ideal prediction performances and compared with the learned models.

We found that with a state dimension equal to that of the true model, RNN PSID achieved ideal prediction for both the behavior and neural signals (**S Fig. 2a,c**). Moreover, even given a minimal state dimension equal to the true number of behaviorally relevant state dimensions, RNN PSID still achieved ideal prediction for behavior (**S Fig. 2b**). These results show that for linear models, similar to the analytical subspace-based PSID from our earlier work^1^, the linear RNN PSID using numerical optimization can also learn behaviorally relevant neural dynamics using low-dimensional latent states. Further, if learning of all neural dynamics regardless of relevance to behavior is of interest, linear RNN PSID can do so given enough state dimensions. We next found that the performance of linear RNN PSID and subspace-based PSID^1^ were similar across various regimes of training samples, with subspace-based PSID^1^ very slightly outperforming linear RNN PSID when learning models with low-dimensional states with very small sample sizes on the order of 1000 (**S Fig. 2b**). Given this similar performance, for simplicity and better consistency hereon after we use linear RNN PSID as our linear modeling method and refer to it simply as linear PSID.

We next validated RNN PSID in numerical simulations with nonlinear models and confirmed that it can be used to successfully dissect the origin of nonlinearity in the data. We simulated random systems where only one of the model parameters were set to a random nonlinear function (Methods). We then used RNN PSID to fit models with the nonlinearities isolated in different parameters and compared them. We found that the RNN PSID model that achieved the best behavior decoding and neural self-prediction was the model that had the nonlinearity in the correct model parameter (**S Fig. 3**), suggesting that RNN PSID could be used to determine the origin of nonlinearity. We will thus use a similar approach in real datasets to dissect nonlinearities and interpret them.

### Nonlinear modeling of behaviorally relevant neural dynamics across four tasks

We used RNN PSID to study the behaviorally relevant neural dynamics in neural data from four monkeys performing four different tasks and having recordings across different brain regions (**Fig 2**, Methods). In the first task, the monkey made naturalistic 3D reach, grasp and return movements while the joint angles in the arm, elbow, wrist, and fingers were tracked as the behavior (**Fig. 2a**)^1,26^. In the second task, the monkey made saccadic eye movements to a randomly selected target out of eight possible targets on a screen, with the 2D eye position tracked as the behavior (**Fig. 2e**)^1,27^. In the third task, which was from a publicly available dataset^28,29^, the monkey made sequential 2D reaches on a screen using a cursor controlled with a manipulandum, while the 2D cursor position and velocity were tracked as the behavior (**Fig. 2i**). In the fourth task, which was from another publicly available dataset^30^, the monkey made 2D reaches to random targets in a grid via a cursor that mirrored the monkey’s fingertip movements, for which the 2D position and velocity were tracked as the behavior (**Fig. 2m**). In tasks 1 and 4, primary motor cortical activity was modeled. For tasks 2 and 3, prefrontal cortex and dorsal premotor cortical activities were modeled, respectively. In all datasets, we took the Gaussian smoothed spike counts as the neural signal to be modeled (Methods). Similar results also held for LFP activity as will be shown in a later section, showing the generality across neural signal modalities.

**Fig. 2.**
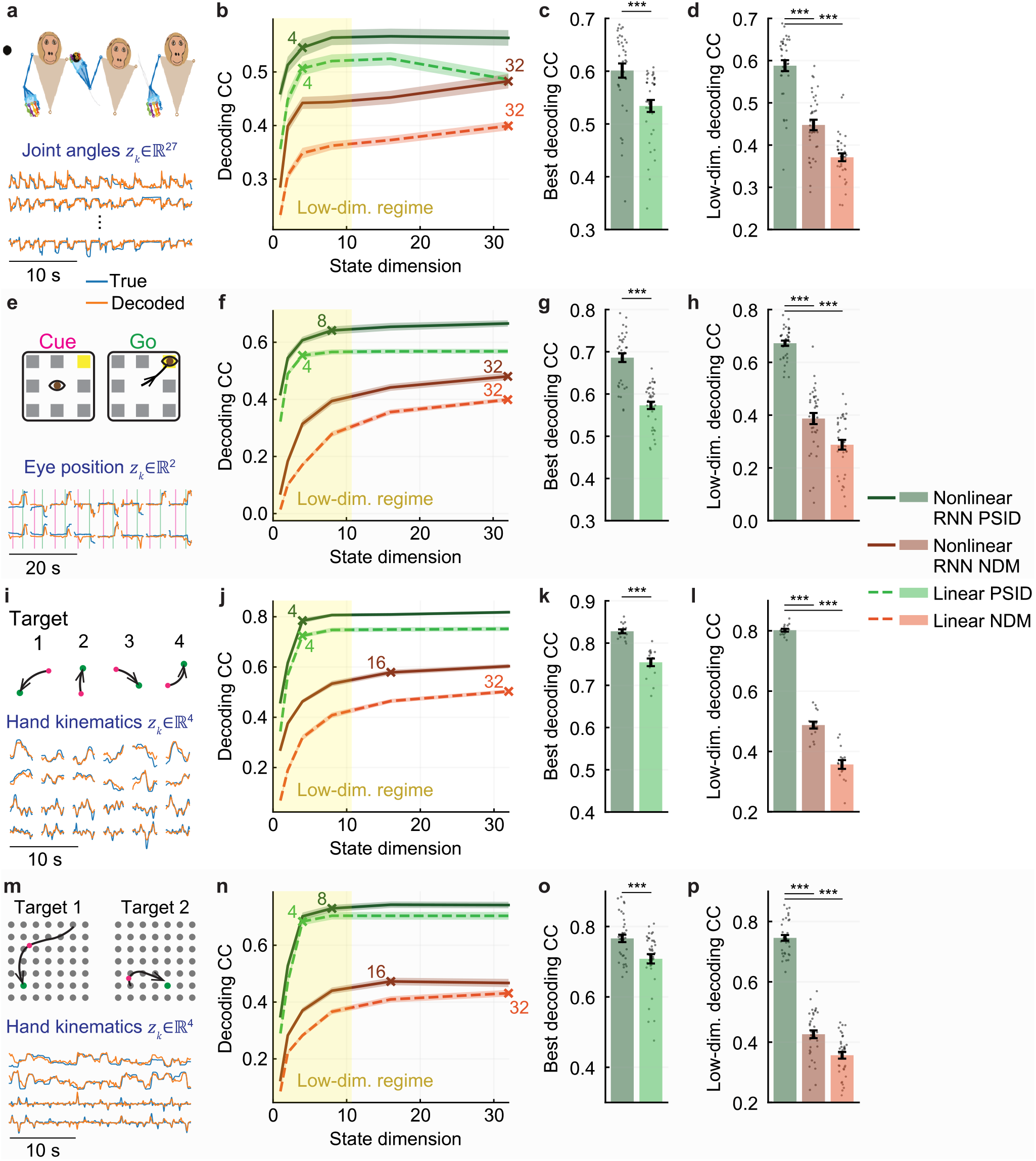
RNN PSID reveals more accurate models of behaviorally relevant neural dynamics by capturing nonlinearity while also being preferential. (**a**) The 3D reach task, along with example true and decoded behavior dimensions, decoded using RNN PSID. (**b**) Cross-validated decoding accuracy (CC) achieved by variations of linear/nonlinear RNN PSID/NDM, for different latent state dimensions. For nonlinear RNN PSID/NDM, the nonlinearities are selected automatically based on the training data to maximize behavior decoding accuracy (flexible nonlinearity; Methods). Solid lines show the average across sessions and folds (*N* = 35) and the shaded areas show the s.e.m. The latent state dimension at which each method reaches the average decoding accuracy that is within 5% of the shown peak decoding accuracy is marked with a cross. (**c**) Peak decoding accuracy achieved by nonlinear and linear RNN PSID, by choosing the state dimension in each session and fold as the smallest that reaches peak decoding. Bars show the mean, whiskers show the s.e.m., and dots show all data points (*N* = 35). Asterisks show significance level for a one-sided Wilcoxon signed-rank test (*: *P* < 0.05, **: *P* < 0.005, and ***: *P* < 0.0005). (**d**) Decoding accuracy of nonlinear RNN PSID versus linear/nonlinear RNN NDM, at the latent state dimension for which RNN PSID reaches within 5% of its peak. Notation is as in (c). (**e-h**) Same as (a-d) for the second dataset, with saccadic eye movements (*N* = 35). (**i**-**l**) Same as (a-d) for the third dataset, with sequential cursor reaches controlled via a 2D manipulandum (*N* = 15). (**m**-**p**) Same as (a-d) for the fourth dataset, with random grid virtual reality cursor reaches controlled via fingertip position (*N* = 35). For all PSID methods, only the first stage was used (i.e. *n*_1_ = *n*_*x*_).

We modeled the neural activity and behavior in each dataset using nonlinear RNN PSID with different latent state dimensions and computed the cross-validated accuracy of causally decoding behavior from past neural activity (**Fig. 2b,f,j,n**). We then compared the peak behavior decoding with that of linear PSID to assess the level of nonlinearity in each dataset (**Fig. 2c,g,k,o**). We found that in all datasets, nonlinear RNN PSID achieved significantly more accurate decoding than linear PSID, suggesting that there is nonlinearity in behaviorally relevant dynamics (**Fig. 2c,g,k,o**). In addition, across all datasets, nonlinear RNN PSID reached within 5% of peak decoding using only 4-8 latent state dimensions (**Fig. 2b,f,j,n**), suggesting that behaviorally relevant neural dynamics are largely low-dimensional, with much smaller dimension than what would be implied with non-preferential NDM methods (16-32). Using the same low latent state dimensions, linear and nonlinear RNN NDM achieved much lower behavior decoding accuracy than nonlinear RNN PSID (**Fig. 2d,h,l,p**), suggesting that neural dynamics are not dominated by behaviorally relevant dynamics and thus these latter dynamics can be missed or confounded if not prioritized during learning. This highlights the capability of RNN PSID as a nonlinear dynamic modeling and dimensionality reduction approach that preferentially preserves behaviorally relevant neural dynamics.

### Overall neural dynamics could also be learned by RNN PSID

The optional second stage of RNN PSID can learn any additional neural dynamics that were not predicted using the latent states from the first stage, which are extracted to be relevant to the specific measured behavior (Methods). Thus, we next used both stages of RNN PSID to also learn other neural dynamics beyond the behaviorally relevant ones learned in the first stage. To evaluate the model, we found the best self-prediction that could be achieved by increasing the latent state dimension and also the corresponding behavior decoding (Methods). A model that has higher accuracy for both behavior decoding and neural self-prediction provides a better explanation of the data overall.

We found that in all datasets, compared with linear/nonlinear RNN NDM or linear PSID, nonlinear RNN PSID reached better behavior decoding accuracy while being as accurate or better in terms of peak neural self-prediction (**Fig. 3**). More specifically, in nonlinear RNN PSID, we use the training data to learn models with various combinations of nonlinearities. Then among these learned models, we can select one model based on training data to be used in the test data (Methods). The criteria for this model selection can be based on achieving either better behavior decoding or better neural self-prediction (Methods). Note that the criteria for selecting among the learned models with different nonlinearities is independent of the internal objective functions used in RNN PSID to learn each model, which are always fixed (**S Fig. 1**; Methods). We found that RNN PSID with nonlinearity selected based on neural self-prediction was better in reaching the best performance frontier and simultaneously achieving accurate behavior decoding and neural self-prediction (**Fig. 3**). Selection of nonlinearity based on behavior decoding resulted in statistically significant improvement in decoding only in one of the datasets (**S Fig. 4h,i**), in which case both selection criteria led to models on the performance frontier (**Fig. 3f**).

**Fig. 3.**
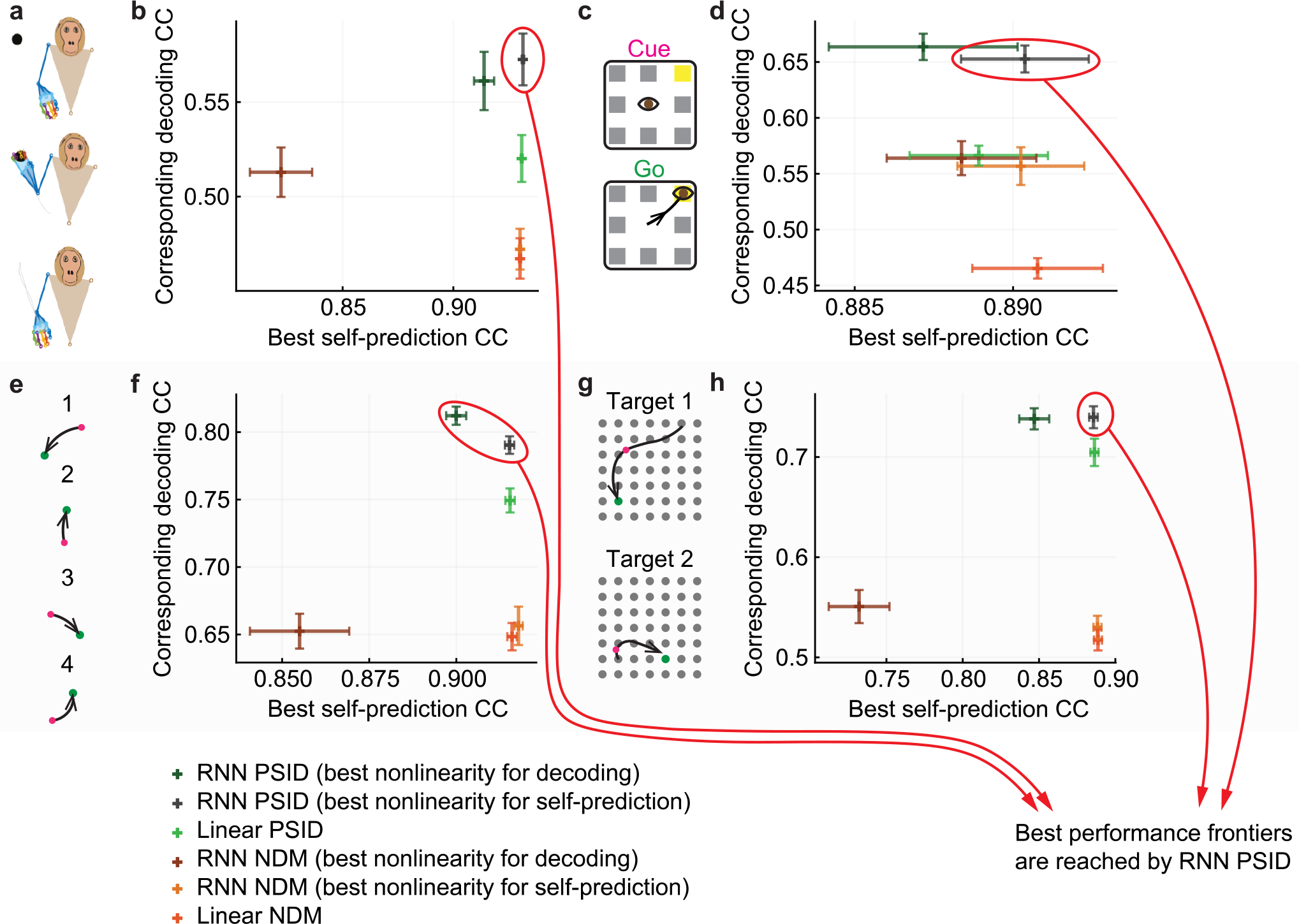
RNN PSID more accurately learns the behaviorally relevant neural dynamics while also capturing the overall neural dynamics as accurately as other methods. (**a**) The 3D reach task. (**b**) Peak neural self-prediction accuracy achieved by each method shown on the horizontal axis versus the corresponding behavior decoding accuracy on the vertical axis. Latent state dimension for each method in each session and fold is chosen as the smallest that reaches peak neural self-prediction. The plus on the plot shows the mean self-prediction and decoding accuracy across sessions and folds (*N* = 35), and the horizontal and vertical whiskers show the s.e.m. for these measures, respectively. Models whose paired (self-prediction, decoding) measure leads to a plus toward the most upper-right corner of the plot lie on the best performance frontier as they have better performance in both measures and thus better explain the data (for definition of frontier see results). (**c**-**d**) Same as (a-b) for the second dataset, with saccadic eye movements (*N* = 35). (**e**-**f**) Same as (a-b) for the third dataset, with sequential cursor reaches controlled via a 2D manipulandum (*N* = 15). (**g**-**h**) Same as (a-b) for the fourth dataset, with random grid virtual reality cursor reaches controlled via fingertip position (*N* = 35). For all RNN PSID variations, the first 16 latent state dimensions are learned using stage 1 and the remaining are learned using stage 2 (i.e. n_1_ = 16). For more details, see **S Figs. 3–4**.

### Neural self-prediction benefited from nonlinear modeling at low dimensions

Comparing nonlinear RNN PSID and nonlinear RNN NDM with their linear counterparts across dimensions, we further found that at low dimensions nonlinearity was needed both for accurate behavior decoding (**S Fig. 4**) and for accurate neural self-prediction (**S Fig. 5**). However, in terms of peak neural self-prediction, given sufficiently high latent state dimensions, linear methods reached similar peak performance to nonlinear methods (**Fig. 3**, and **S Fig. 5**). In contrast, in terms of behavior decoding, nonlinear RNN PSID significantly outperformed linear methods even at high dimensions (**Fig. 3** and **S Fig. 4**). These results suggest that while there likely exist some nonlinear dynamics in neural activity, with high enough latent state dimensions, linear models can become good approximations of the overall neural dynamics as long as performing neural self-prediction alone is of interest. However, even with such high latent state dimensions, linear models cannot achieve as accurate of a description for behaviorally relevant neural dynamics as evident from their inferior peak behavior decoding. To further understand the nature of this nonlinearity, we next investigated the model parameters whose nonlinearity was most essential for describing these behaviorally relevant neural dynamics.

### Nonlinear behaviorally relevant neural dynamics could largely be explained with a nonlinear behavior readout model parameter

Given the flexibility of RNN PSID in selectively allowing nonlinearity in each model parameter, we next used it to dissect nonlinearities and investigate the origin of nonlinearity in each dataset. To do so, we used the same hypothesis testing procedure that was validated in our simulations to correctly find the origin of nonlinearity (**S Fig. 3**). We built alternative models with different individual model parameters being nonlinear and compared the resulting models in terms of behavior decoding and neural self-prediction to each other and to fully nonlinear models that could have nonlinearity in all or any combination of parameters (flexible nonlinearity; Methods). Strikingly, we found that having nonlinearity only in the behavior readout parameter *C*_*z*_ was sufficient for achieving high decoding and self-prediction accuracy across all datasets despite their diverse behavioral tasks and neural recordings (**Fig. 4**). First, models with nonlinearity only in the behavior readout parameter *C*_*z*_ reached the best behavior decoding accuracy compared with models with other individual nonlinearities (**Fig. 4b,f,j,n**), while reaching almost the same decoding accuracy as fully nonlinear models with flexible selection of nonlinearity in all or any combination of parameters (**Fig. 4b,f,j,n**). Second, in terms of reaching the best neural self-prediction, again models with nonlinearity only in the behavior readout reached a peak self-prediction accuracy that was unmatched by other types of individual nonlinearity (**Fig. 4c,g,k,o**). Taken together, models with nonlinearity in the behavior readout parameter *C*_*z*_ achieved the best behavior decoding accuracy, while simultaneously matching the best of all other linear or nonlinear models in terms of neural self-prediction (**Fig. 4d,h,l,p**). These results suggests that across all four datasets with their diverse array of behavioral tasks, the origin of nonlinearity is not necessarily in the dynamics of low-dimensional states themselves, but could be mostly confined to how these neural dynamics are mapped to behavior (i.e. in the behavior readout).

**Fig. 4.**
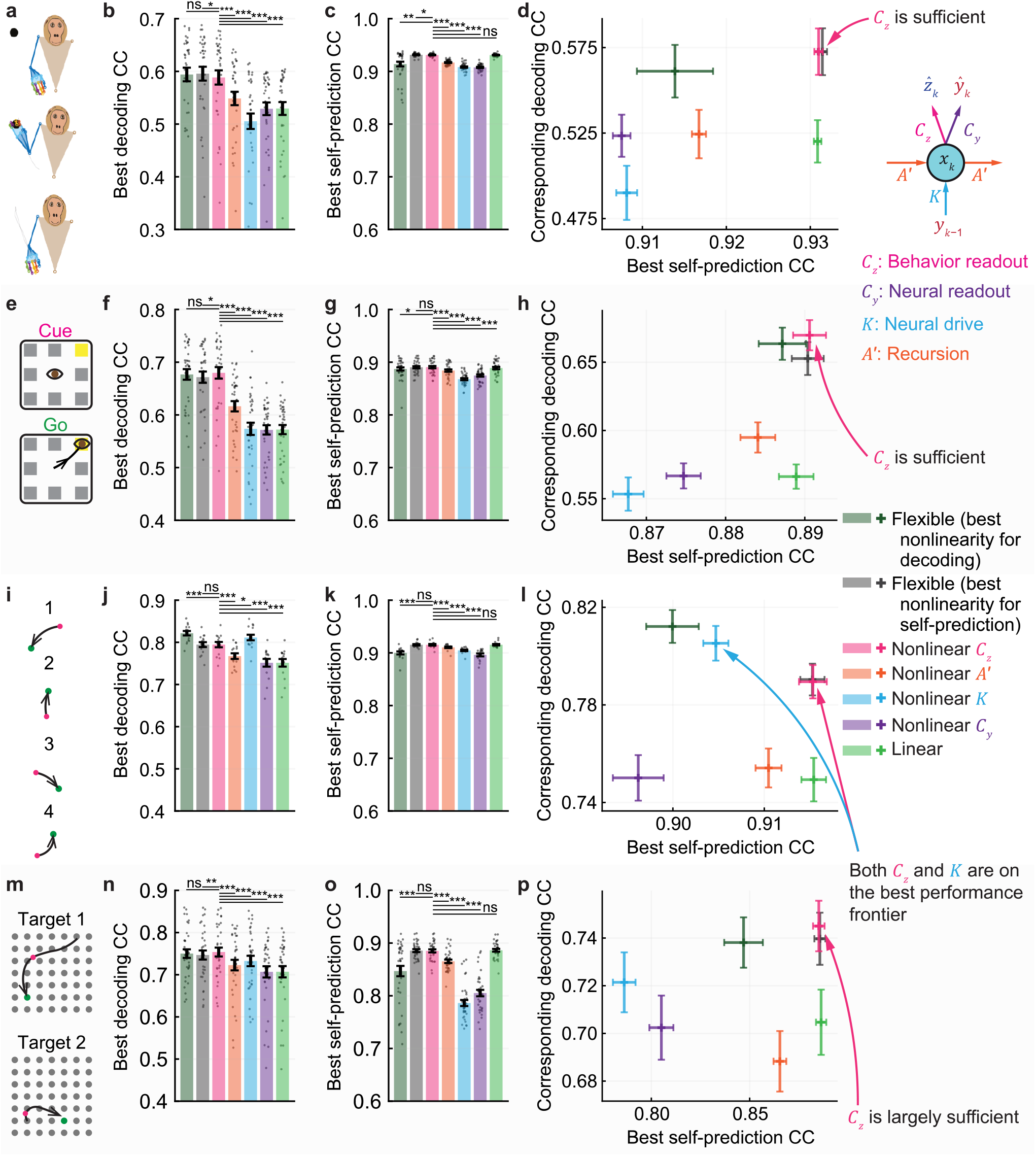
RNN PSID reveals that across datasets, nonlinearities can be largely captured by the behavior readout of the model. (**a**) The 3D reach task. (**b**) Peak cross-validated decoding accuracy (CC) achieved by variations of RNN PSID with different nonlinearities. RNN PSID variations that have only one nonlinear parameter (e.g. *C*_*z*_) use a fixed network structure with one 64-unit hidden layer for that parameter and keep all other parameters linear (Methods). Linear and flexible nonlinear results are as in **S Fig. 4**. State dimension in each session and fold is chosen as the smallest that reaches peak decoding. Bars, whiskers, and dots are defined as in **Fig. 2b** (*N* = 35). (**c**) Same as (b), showing the peak cross-validated neural self-prediction accuracy (CC) for each nonlinearity (linear and flexible nonlinear results are as in **S Fig. 5**). (**d**) The corresponding decoding accuracy for each peak neural self-prediction (linear and flexible nonlinear results are as in **Fig. 3**). Notation is as in **Fig. 3**. (**e**-**h**) Same as (a-d) for the second dataset, with saccadic eye movements (*N* = 35). (**i**-**l**) Same as (a-d) for the third dataset, with sequential cursor reaches controlled via a 2D manipulandum (*N* = 15). (**m**-**p**) Same as (a-d) for the fourth dataset, with random grid virtual reality cursor reaches controlled via fingertip position (*N* = 35). For all RNN PSID variations, the first 16 latent state dimensions are learned using stage 1 and the remaining are learned using stage 2 (i.e. n_1_ = 16).

Given these results, we next investigated whether an end-to-end learning of a dynamic model that only has nonlinearity in the behavior readout parameter *C*_*z*_ has any benefits compared with a linear model that is upgraded in a post-hoc step to have a nonlinear behavior readout. To compare these cases, we took the already-learned latent states from linear PSID models, learned a nonlinear mapping from these latent states to behavior, and replaced behavior readout of the linear models with this nonlinear mapping that was learned post-hoc. We used the exact same neural network architecture as was used for the behavior readout parameter *C*_*z*_ in the previous results, so the final model had identical structure to the model that was learned end-to-end, with their only difference being in their learned network weights. We found that in all datasets, the final behavior decoding accuracy from the post-hoc learning was significantly worse than the accuracy from the end-to-end learning of a model with nonlinear behavior readout parameter *C*_*z*_ (*P* ≤ 0.0027, one-side signed-rank, *N* ≥ 15). This is because despite having the same architecture, the model that is initially fully linear would extract different latent states that are learned based on the assumption of a linear behavior readout, so they may not find and fully exploit the low-dimensional neural subspaces that are informative of behavior with a nonlinear behavior readout. These results highlight the importance of RNN PSID being able to directly learn models with nonlinearities in arbitrary sets of parameters, even in datasets where nonlinearity in only the behavior readout may be sufficient for accurate modeling.

### Similar results held for raw LFP and LFP band power activity, with raw LFP activity showing the largest gain from nonlinear modeling

In three of the four tasks, LFP activity was also available. We thus performed similar analyses for two types of features from LFP activity. First, we used the raw LFP activity, downsampled to the sampling rate of the behavior (i.e. 50ms time steps), as the neural feature (Methods). In the context of the motor cortex, downsampled raw LFP is also referred to as the local motor potential (LMP)^31–33^, and has previously been used to decode behavior^1,31–34^. Second, we extracted signal power in standard frequency bands from delta (0.1-4 Hz) to high gamma (130-170 Hz) as the neural feature (Methods). We found that across all tasks and for both types of LFP features, RNN PSID more accurately learned behaviorally relevant neural dynamics than linear PSID or non-preferential RNN NDM (**S Figs. 6–7**), while also achieving similar or higher accuracy in neural self-prediction (**S Figs. 6–7**). Moreover, interestingly, for both LFP features, having nonlinearity only in the behavior readout parameter was again largely sufficient for learning neural dynamics (**S Figs. 8–9**). These results suggest that our earlier results on neural population spiking activity generalize to dynamics in other neural modalities such as raw LFP activity and LFP band power activity.

We next compared the nonlinear RNN PSID and linear PSID results across neural modalities to quantify the amount of nonlinearity in each neural modality (**Fig. 5**). We found that in all three tasks, raw LFP activity had the highest gain in behavior decoding accuracy by going from linear to nonlinear modeling. Notably, when using RNN PSID to find the best nonlinear modeling and transformation, raw LFP activity reached more accurate behavior decoding accuracy than LFP band powers in all tasks (**Fig. 5b,e,h**). In the task with saccadic eye movements, raw LFP activity even significantly surpassed the decoding accuracy of spiking activity (**Fig. 5e**). These results suggest that equipped with a model that can learn flexible nonlinear dynamics, behaviorally relevant information that exists in LFP activity but may normally be missed can be more accurately learned. Further, computing LFP powers involves a pre-specified nonlinear transformation on raw LFP. As such, these results show that nonlinear RNN PSID automatically finds the dynamical nonlinear transformation on raw LFP that does better in explaining neural/behavior data than the transformation to compute these pre-specified power band features.

**Fig. 5.**
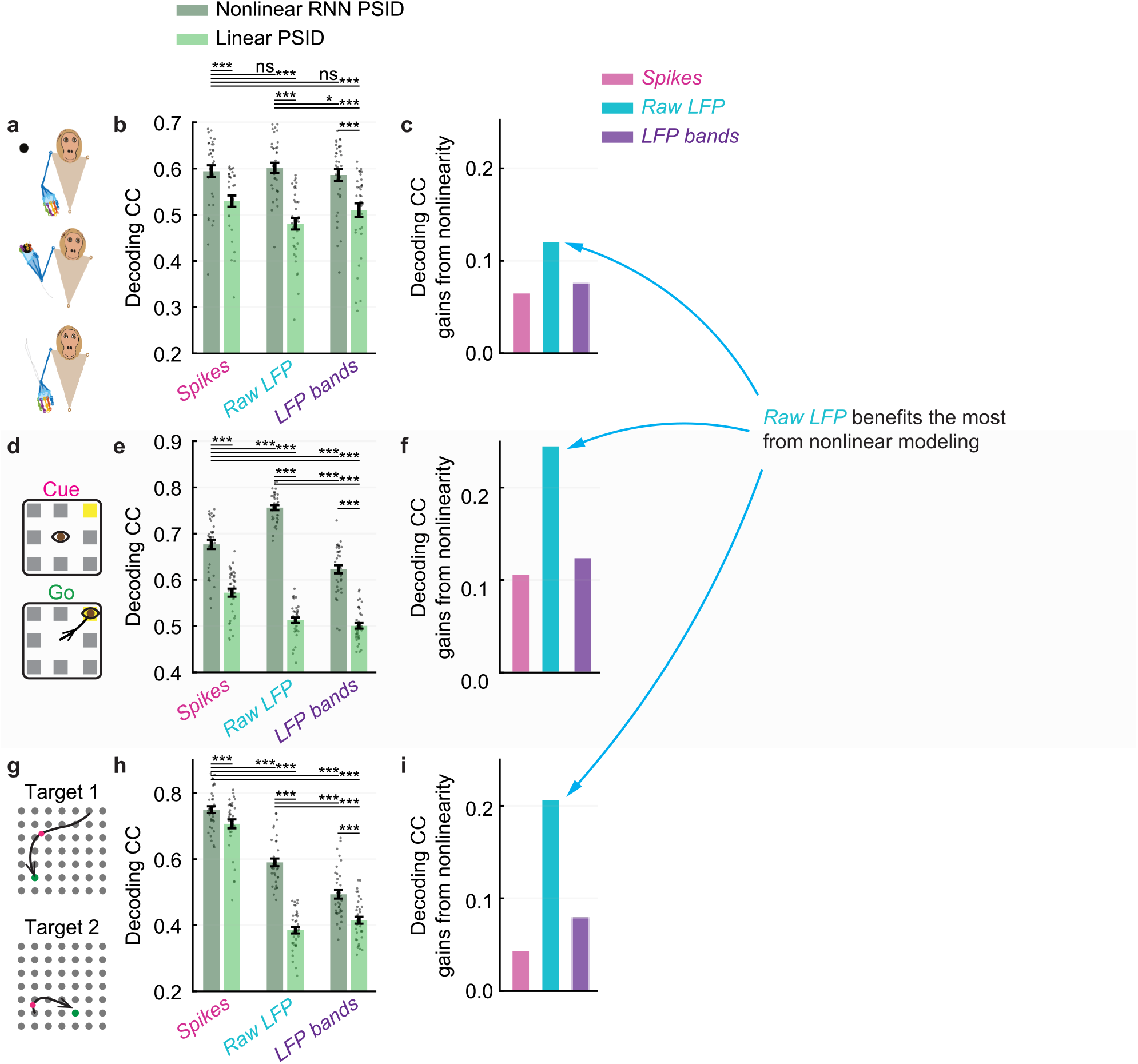
Raw LFP activity benefits the most from nonlinear modeling, compared with LFP bands or spiking activity. (**a**) The 3D reach task. (**b**) Peak cross-validated decoding accuracy (CC) achieved by variations of nonlinear RNN PSID and linear PSID. Results are shown for spiking activity, raw LFP activity, and LFP band power activity (Methods), as also shown in **Fig. 3** and **S Figs. 6–7**. For nonlinear RNN PSID, the nonlinearity is selected automatically based on the training data to maximize behavior decoding accuracy (Methods). Bars, whiskers, and dots are defined as in **Fig. 2c**. (**c**) The difference between the nonlinear and linear results from (b). When doing flexible nonlinear modeling with RNN PSID, raw LFP activity achieves better decoding than the LFP power bands that are pre-specified nonlinear transformations of a more wideband version of this raw activity (Methods). (**d**-**f**) Same as (a-c) for the second dataset, with saccadic eye movements. (**g**-**i**) Same as (a-c) for the fourth dataset, with random grid virtual reality cursor reaches controlled via fingertip position.

### Preferential modeling of behaviorally relevant neural dynamics uncovered distinct low dimensional representations for these dynamics

Given that RNN PSID can prioritize learning of behaviorally relevant neural dynamics, it can be used to perform dimensionality reduction while preserving these dynamics. To demonstrate this, we learned models with 2-dimensional latent states and visualized their latent state trajectory during different epochs and conditions of each task. In each dataset, we averaged the latent states across trials with similar movement conditions (**Fig. 6d,g,j**, Methods) and then compared the obtained average trajectories across movement conditions. We found that consistently, RNN PSID extracted latent states from neural activity that were clearly different depending on the behavior condition (**Fig. 6b,e,h,k**), whereas RNN NDM extracted latent states that did not as clearly dissociate different conditions (**Fig. 6c,f,i,l**). Moreover, quantitively, states extracted using RNN PSID could more accurately decode behavior than those extracted using RNN NDM (see **Fig. 2b,f,j,n** for state dimension of 2). Notably, for the first dataset, in our earlier work^1^, we had compared the latent trajectories for subspace-based linear PSID versus linear NDM and had found that PSID shows distinct reverse-rotational patterns across reach and return movement conditions, as is also the case here for RNN PSID versus RNN NDM (**Fig. 2b-c**). These results thus complement our prior work^1^ by showing that even nonlinear NDM models are not able to uncover the distinct behaviorally relevant dynamics that reverse their rotation direction in this dataset. Together, these results show that even including nonlinearity in NDM methods cannot help uncover these dissociated behaviorally relevant states but that extracting these states critically requires the preferential element of RNN PSID. Moreover, these results suggest RNN PSID uncovers distinct low-dimensional neural dynamics that are more behaviorally relevant but may be missed by even nonlinear NDM methods, which do not prioritize behaviorally relevant neural dynamics.

**Fig. 6.**
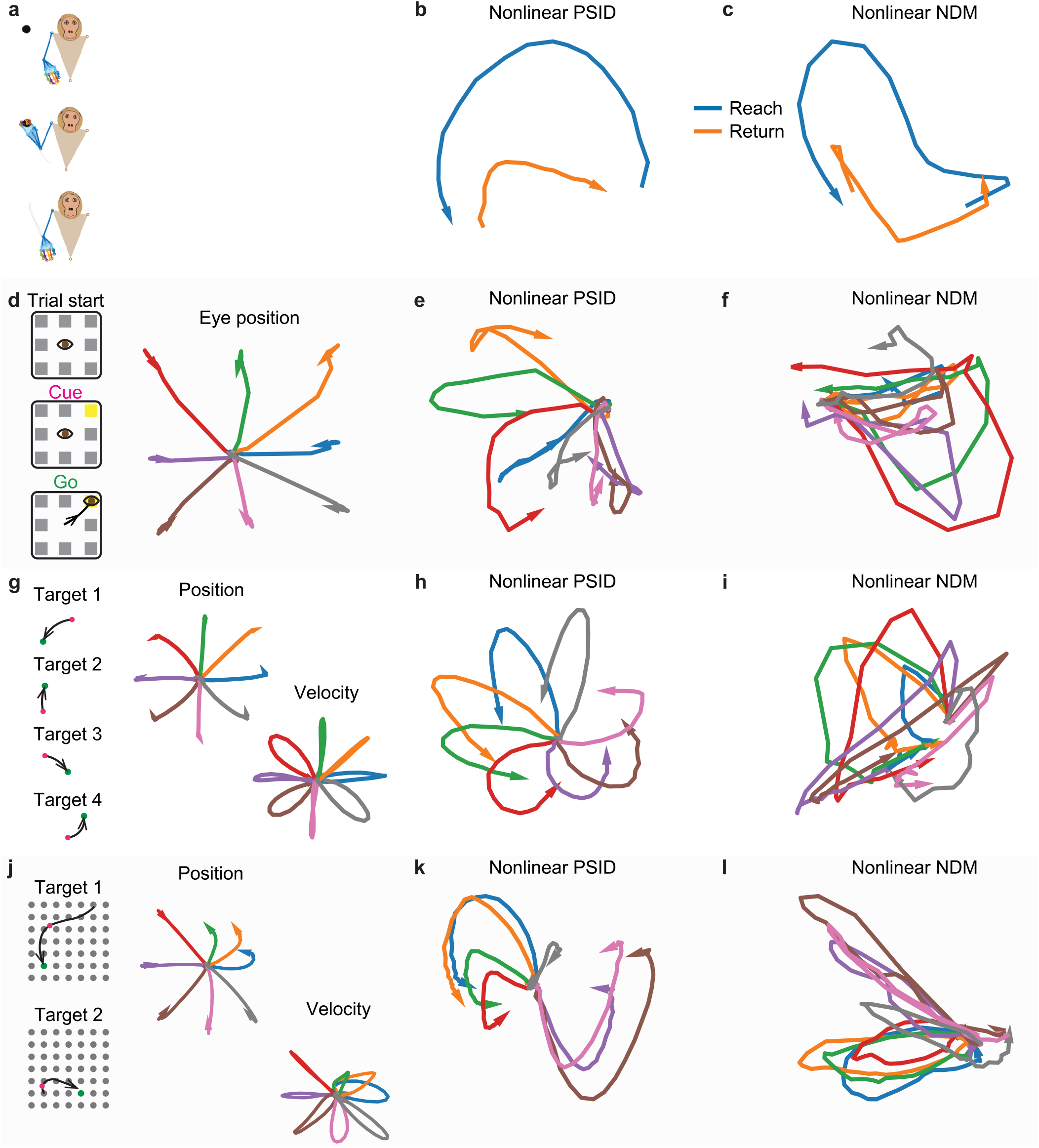
Nonlinear RNN PSID extracted distinct low dimensional latent states for all datasets, which were more behaviorally relevant than those extracted using nonlinear RNN NDM. (**a**) The 3D reach task. (**b**) The latent state trajectory for 2D states extracted from spiking activity using nonlinear RNN PSID, averaged across all reach and return epochs across sessions and folds. RNN PSID revealed latent states with rotational dynamics that reversed direction during reach versus return epochs, which is consistent with the behavior roughly reversing direction. (**c**) Same as (b), for 2D states extracted using nonlinear RNN NDM. In contrast to RNN PSID, latent state extracted using RNN NDM showed rotational dynamics that did not reverse direction during reach versus return periods, thus were less congruent with behavior. (**d**) Saccadic eye movement task. Trials are averaged depending on the eye movement direction. (**e**) The latent state trajectory for 2D states extracted using RNN PSID, averaged across all trials of the same movement direction condition across sessions and folds. (**e**) Same as (d), for 2D states extracted using nonlinear NDM. (**g-i**) Same as (d-f), for the third dataset, with sequential cursor reaches controlled via a 2D manipulandum. (**j**-**l**) Same as (d-f) for the fourth dataset, with random grid virtual reality cursor reaches controlled via fingertip position. Overall, latent states extracted by RNN PSID were clearly different depending on the behavior condition (c,e,h,k), whereas RNN NDM extracted latent states did not as clearly dissociate different conditions (c,f,I,l).

## Discussion

Here, we developed a new modeling approach termed RNN PSID that enables nonlinear dynamic modeling with causal decoding, can dissect the origin of nonlinearity through its flexible nonlinearity, and can dissociate and prioritize behaviorally relevant neural dynamics from other neural dynamics. These attributes make RNN PSID uniquely suitable for investigations of the nature of nonlinearity in the dynamical neural encoding of behavior across diverse domains of neuroscience, and for developing real-time brain-machine interfaces (BMIs) across applications.

Here, by comparing various types of nonlinearities, we found that models with only nonlinearity in the behavior readout best described behaviorally relevant and overall neural dynamics across all four distinct tasks. We found this by devising a hypothesis testing procedure that was enabled via the flexible nonlinearity of RNN PSID, and by first validating this procedure within simulations with known origins for nonlinearity (**Fig. 1b**). This is an interesting finding since as demonstrated in the simulations (**Fig. 1b**), technically the origin of nonlinearity in RNN PSID could also be in any one (or more) of the following four options (**Fig. 1a**): the neural drive (mapping from neural activity to the low-dimensional subspace), the recurrent dynamics (mapping from latent state from one time step to the next), and the neural or behavioral readout (mapping from latent state to neural or behavioral observations). The fact that of these four types of nonlinearities, a nonlinearity in behavior readout is largely sufficient for explaining the data across tasks suggests the following interesting possibility: that across all four tasks, neural dynamics in the recorded brain areas are largely describable with linear dynamics (given large enough latent state dimensions), but additional nonlinearities are introduced somewhere along the neuromuscular pathway that goes from the recorded area to the measured behavior. This is also consistent with our finding that given enough latent state dimensions, linear models are sufficient for accurate neural self-prediction, but they still fail to reach behavior decoding that is as accurate as RNN PSID with a nonlinear behavior readout (**Fig. 4d,l,p**).

It is important to note that in general, in some cases it may be mathematically possible to equivalently explain the same neural or behavior data with multiple types of nonlinearities (e.g. either with a nonlinear neural drive, or a nonlinear readout). However, in numerical simulations, we find that when both neural and behavior data are considered together, usually only the correct type of nonlinearity explains the data best (**S Fig. 3**). It is also possible that the best model describing the data requires two or more of the four parameters to be nonlinear. But in our datasets, models with nonlinearity only in behavior readout were always on the performance frontier and could not be considerably outperformed by models with more than one nonlinearity in terms of both the decoding and neural self-prediction measures (**Fig. 4**). Only in one dataset a flexible search over nonlinearities achieved significantly better decoding from population spiking activity than having the nonlinearity only in the behavior readout (**Fig. 4j**), but even in that dataset the latter explained the overall neural dynamics more accurately (**Fig. 4j**).

We found similar results for three neural modalities: spiking activity, LFP band power activity, and raw LFP activity. For all three data types, RNN PSID more accurately learned behaviorally relevant neural dynamics compared with linear PSID and non-preferential RNN NDM as reflected in its better decoding, while also achieving similar or more accurate neural self-prediction. Moreover, for all three data types, having nonlinearity only in the behavior readout was largely sufficient to explain the data. These results demonstrate the general applicability of RNN PSID to both spiking and LFP activity and its general utility in testing hypotheses about the origin of nonlinearity. Notably, the raw LFP activity benefited the most from nonlinear modeling using RNN PSID and outperformed LFP powers in all tasks in terms of decoding, suggesting that equipped with an automatic learning of nonlinear models as enabled by RNN PSID, the benefit of manually extracting traditionally used features such as LFP band powers may be limited and better nonlinear mappings may be automatically learned.

An important challenge in nonlinear dynamic modeling is building a decoder that can take neural observations and produce their latent representation and predict behavior. Here, we directly learn the model in predictor form such that after learning, the decoder is readily available, rather than requiring expensive computations such as an iterative expectation maximization^3,13^. Moreover, decoding is done using an RNN, without the need for computationally expensive sampling and averaging that is needed by variational sequential autoencoders^2^. More importantly, unlike sequential autoencoders, the decoding is fully causal and uses only past neural information, rather than using future data that would not be recorded yet. These characteristics make RNN PSID an attractive model for real-time closed-loop BMI applications where fast feedback is necessary to maintain natural interaction with the BMI^35^.

Recent work has used sequential autoencoder RNNs to smooth single-trial neural activity via nonlinear dynamic modeling^2^. However, this approach does not consider behavior during the learning of the dynamics, and thus does not dissociate or prioritize the learning of behaviorally relevant neural dynamics resulting in their potentially less accurate learning^1^. Other prior works have used RNNs for causal decoding of behavior from neural activity^15,16^. These works have similarities to the first optimization in the first stage of RNN PSID in that they optimize the behavior prediction, but they do not learn the mapping from the RNN latent states to neural activity, which is done using a second optimization in the first stage of RNN PSID to enable neural self-prediction using the behaviorally relevant dynamics (**S Fig. 1**). In addition, unlike what the second stage of RNN PSID enables, these prior works do not model additional neural dynamics beyond those that decode behavior, and thus do not aim to dissociate the two types of neural dynamics. Moreover, each of these works^15,16^, including those with non-causal sequential autoencoders^2^, use specific nonlinear RNN structures whereas in RNN PSID the nonlinear structure is automatically selected in a way that best suits the training data within an inner cross-validation (Methods). Finally, importantly, here unlike prior works, we further dissect the nonlinearity and explore how one can isolate the nonlinearity in specific parameters of the nonlinear RNN, each of which has interpretable roles. These new advances make RNN PSID a comprehensive framework for understanding nonlinear neural dynamics that give rise to behavior.

Beyond enabling prioritized learning of behaviorally relevant neural dynamics with support for nonlinear dynamics, RNN PSID also supports several other applications by incorporating new innovations that we will fully present in our future work^19^. First, given the numerical optimization approach, RNN PSID is also applicable if behavior or neural signals are intermittently sampled, such as when modeling sparsely measured behaviors such as mood questionnaires^36^, which will be needed toward understanding neuropsychiatric conditions and developing new therapies for them^12,37^. Second, RNN PSID supports modeling neural activity with non-Gaussian (e.g. Poisson) distributions, which could benefit the modeling of some data types such as spiking activity sampled at millisecond bins and lead to better closed-loop BMI performance^35,38^. Third, RNN PSID can support modeling behavior with non-Gaussian (e.g. categorical) distributions, such as decision choices. These additional capabilities will make RNN PSID of potential benefit in various additional neuroscience and neurotechnology applications as we will show in our future work^19^.

Taken together, RNN PSID is a general new approach for preferentially modeling behaviorally relevant neural dynamics that supports nonlinear dynamics, enables flexible exploration of their origin while also allowing for causal decoding of latent dynamics and behavior.

## Methods

### Model formulation

The model used by RNN PSID is provided in equation (1). We write this model as a nonlinear generalization of the predictor form^21,39^ of the linear models such that we can enable flexible and interpretable dissection of nonlinearities and causal prediction. For a full description of the motivation for the model formulation and its relation to linear models in predictor form please refer to **S Note 1**. Equation (1) corresponds to an RNN computation graph (**Fig. 1b**) and thus its parameters can be learned using standard tools for numerical optimization^20^. For this RNN, neural activity *y*_*k*_ constitutes the input and predictions of neural and behavioral signals are the outputs (**Fig. 1b**) given by

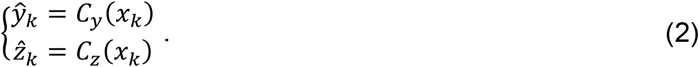

We learn the model parameters by minimizing the mean squared error in the predictions of neural and behavioral signals, which constitute our neural and behavioral losses defined as

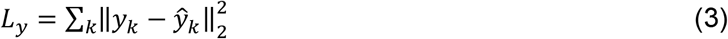

and

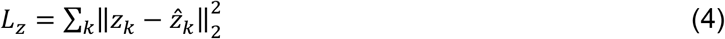

where the sum is over all available samples of *y*_*k*_ and *z*_*k*_, respectively. Importantly, since a key goal of RNN PSID is to dissociate and prioritize the learning of the behaviorally relevant neural dynamics, we devise a two-stage approach for optimizing the above objective functions such that the first stage can prioritize the learning of behaviorally relevant dynamics and the second stage can then learn additional dynamics. We will explain this learning approach in the next section. Once the model parameters are learned, the extraction of latent states *x*_*k*_ involves iteratively applying the first line from equation (1), and predicting behavior or neural activity involves applying equation (2) to the extracted *x*_*k*_.

A more general formulation for the optimization objective in both stages of RNN PSID can be written as

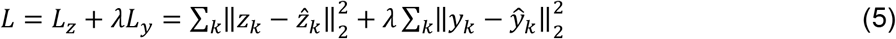

where *λ* determines how much attention is paid to each of the two terms. Here, we use the extreme cases of *λ* = 0 and λ → ∞ for extraction of latent states in stages one and two, respectively. More generally, if dissociation of dynamics is not of interest, one could modify stage one to also incorporate some dynamics beyond the behaviorally relevant dynamics by using a non-zero λ.

### Learning: Two-stage numerical optimization approach

To enable dissociation and prioritization of the behaviorally relevant neural dynamics, we devise a novel two-stage procedure for training the RNN and extracting the latent states. This approach is similar in spirit to our two-stage approach in prior work on dissociating these dynamics for linear dynamic models using subspace-PSID, but is distinct in that subspace-PSID is based on analytical linear algebraic projections^1^ unlike the RNN approach here that is for nonlinear models. The two-stage RNN training approach aims to prioritize the extraction and learning of the behaviorally relevant dynamics in the first stage while also enabling the learning of the rest of the neural dynamics in the second stage such that the latter do not mask or confound the former. We separate the latent states into behaviorally relevant and irrelevant parts (i.e. 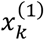 and 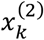), which will be learned in the first and second stages of RNN PSID, respectively. Given state dimension hyperparameters *n*_1_ and *n*_*x*_, we separate the latent states into two parts as

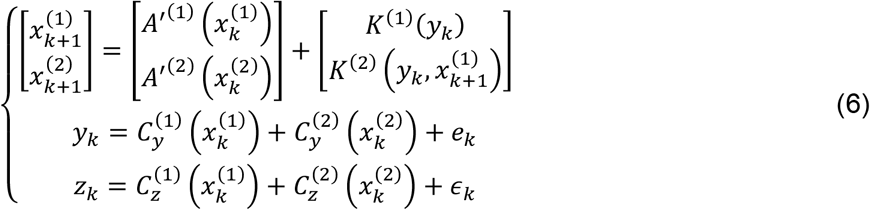

where 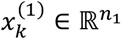 denotes the latent states to be extracted in the first stage and 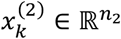, with *n*_2_ = *n*_*x*_ − *n*_1_, denotes those to be extracted in the second stage. The computation graph for equation (6) is provided in **S Fig. 1**. Note that the recursions for computing 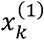 are not dependent on 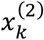, thus allowing the former to be computed without the latter. In contrast, 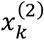 can depend on 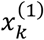 and this dependence is modeled via *K*^(2)^ (**S Note 1**). Note that such dependence of 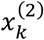 on 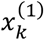 via *K*^(2)^ does not introduce new dynamics to 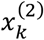 because it doesn’t involve the recursion parameter *A*′^(2)^, which describes the dynamics of 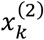.

In the first stage, the objective is to learn behaviorally relevant latent states 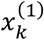 and their associated parameters. This stage consists of two numerical optimizations. In the first optimization, we learn the parameters *A*′^(1)^, 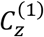, and *K*^(1)^ of the following RNN

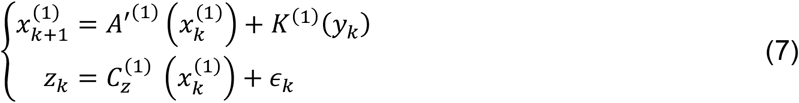

and estimate its latent state 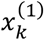 while minimizing the behavior loss *L*_*z*_ (equation (4)). The second optimization uses the extracted latent state 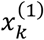 from the RNN and fits the parameters 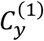 in

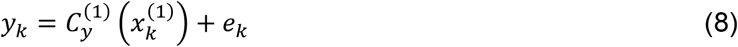

while minimizing the neural loss *L*_*y*_ (equation (3)). Equation (8) can be thought of as a regression model from 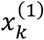 to neural activity *y*_*k*_. This stage concludes the extraction and modeling of behaviorally relevant latent states 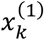.

In the second stage, the objective is to learn any additional dynamics in neural activity that were not learned in the first stage, i.e. 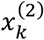 and its associated parameters. To do so, we learn the parameters *A*′^(2)^, 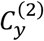, and *K*^(2)^ of the following RNN

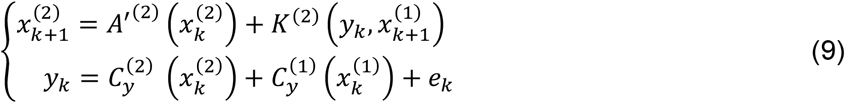

while minimizing the neural loss *L*_*y*_ (equation (3)). Note that in this RNN, *y*_*k*_ and 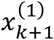 jointly have the role of input and 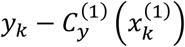 (with the previously learned parameter 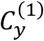) is the output. If we assume the second set of states 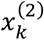 do not contain any information about behavior, we could stop the modeling. However, this may not be the case if the dimension of the states extracted in the first stage (i.e. *n*_1_) is selected to be very small such that some behaviorally relevant neural dynamics are not learned in the first stage. To be robust to such selections of *n*_1_, we can use another numerical optimization to determine based on the data whether and how 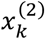 should affect behavior prediction. Thus, a second optimization in the second stage uses the extracted latent state in both stages and fits *C*_*z*_ in

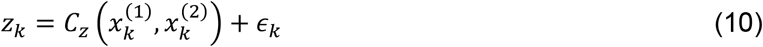

while minimizing the behavior loss *L*_*z*_ (equation (4)). This parameter will replace both 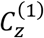 and 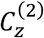 in equation (6). This concludes the learning of all model parameters in equation (6). In this work, when both stages are used together, we do not perform the additional optimization in equation (10), and the prediction of behavior is done solely using the 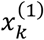 states extracted in the first stage. When only the second stage is used (i.e. *n*_1_ = 0), which we refer to as RNN NDM, the optimization to learn *C*_*z*_ in the second stage is essential and is how the mapping from the latent states to behavior is learned in the RNN NDM case (note in this case we simply have a unified state *x*_*k*_ as there is no dissociation involved).

Finally, the first line of equations (7) and (9), can also be written more generally as

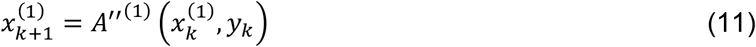

and

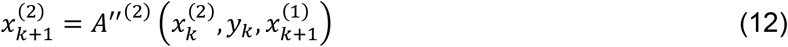

where instead of an additive relation between the two terms of the right-hand side, both terms are combined in nonlinear functions *A*′′^(1)^ and *A*′′^(2)^, which as a special case can still learn the additive relation in equations (7) and (9). Whenever both *A* and *K* are specified to be nonlinear, we use the more general architecture in equations (11) and (12), and if any one of *A* or *K* or both are linear, we use equations (7) and (9).

Once the learning is complete, we also compute the covariances of the neural and behavior residual time series *e*_*k*_ and ∈_*k*_ as Σ_*e*_ and Σ_∈_, respectively. This allows the learned model to be usable in applications where generating new simulated data from the model is desired. This application is not discussed in this work, but an explanation of how new simulated data can be generated form the model in equation (1) is provided in the method section on numerical simulations.

### Behavior decoding and neural self-prediction evaluation metrics

To evaluate the learning, we perform a cross-validation with 5 folds (unless otherwise noted). Specifically, we cut the data from the recording session into five equal segments, leave these segments out one by one as the test data, and train the model only using the data in the remaining segments. Once the model is trained using the neural and behavior training data, we pass the neural test data to the model to get the latent states during the test data using the first line of equation (1). We then pass the extracted latent states to equation (2) to get the one-step ahead prediction of the behavior and neural test data, which we refer to as behavior decoding and neural self-prediction, respectively. Note that only past neural data is used to get behavior and neural predictions and the behavior test data is never used. Given the predicted behavior and neural time series in the data, we then compute the correlation coefficient (CC) between each dimension of these time series and the actual behavior and neural test data. We then take the mean of CC across dimensions of behavior and neural data to get one final cross-validated CC value for behavior decoding and one final CC value for neural self-prediction in this cross-validation fold. After all cross-validation folds are performed, we will have one accuracy value for each fold.

To find the peak behavior decoding (**Fig. 2c,g,k,o**, **Fig. 4b,f,j,n** and **S Fig. 4**) or neural self-prediction (**Fig. 4c,g,k,o** and **S Fig. 5**) accuracy in a given cross-validation fold, we fit models with various latent state dimensions with *n*_*x*_ being different powers of 2 from 1 to 128, and then pick the latent state dimension that achieves the best behavior decoding or neural self-prediction, respectively. Ideally, we want to select a model that simultaneously explains behavior and neural data well as a more accurate representation of both the neural and the behavior data. To do so, given that state dimensions required for reaching peak neural self-prediction are typically higher than the state dimensions required for reaching peak behavior decoding, we find the state dimension that reaches peak neural self-prediction, and report both the neural self-prediction and the corresponding behavior decoding accuracy for the same model (**Fig. 3** and **Fig. 4d,h,l,p**).

### RNN PSID with flexible nonlinearity: automatic determination of appropriate nonlinearity

Each parameter in the RNN PSID model represents an operation in the computation graph of RNN PSID (**Fig. 1b, S Fig. 1**). We solve the numerical optimizations involved in model learning via stochastic gradient descent^20^, which remains applicable for any modification of the computation graph that remains acyclic. Thus, the operation associated with each model parameter (e.g. *A*′, *K*, *C*_*y*_, and *C*_*z*_), can be replaced with any multi-layer neural networks with arbitrarily number of hidden units and layers and the model remains trainable with the same approach. Of course, given that the training data is finite, the typical trade-off between model capacity and generalization error remains^40^. Given that neural networks can approximate any continuous function (with a compact domain)^41^, replacing model parameters with neural networks should have the capacity to learn any nonlinear function in their place. In this work, we use multi-layer feed-forward networks with 1 or 2 hidden layers, each with 64 or 128 units. For all hidden layers we always use a rectified linear unit (ReLU) nonlinear activation (**Fig. 1e**).

We devise a procedure for automatically determining the most suitable combination of nonlinearities for the data, which we refer to as RNN PSID with flexible nonlinearity. In this procedure, for each cross-validation fold in each recording session of each dataset, we try a series of nonlinearities within the training data and select one based on an inner cross-validation within the training data. Specifically, we consider the following options for the nonlinearity. First, each of the four main parameters (i.e. *A*′, *K*, *C*_*y*_, and *C*_*z*_) can be linear or nonlinear, resulting in 16 cases (i.e. 2^4^). In the cases with nonlinearity, we consider four network structures for the parameters, i.e. having 1 or 2 hidden layers and having 64 or 128 units in each hidden layer (**Fig. 1e**), resulting in 61 cases (i.e. 15 × 4 + 1, where 1 is for the fully linear model) overall. Finally, specifically for the recursion parameter *A*′, we also consider modeling it as an LSTM, with the other parameters still having the same nonlinearity options as before, resulting in another 29 cases for when this LSTM recursion is used (i.e. 7 × 4 + 1, where 1 is for the case where the other 3 model parameters are all linear), bringing the total number of considered cases to 90. For each of these 90 considered linear or nonlinear architectures, we perform a two-fold inner cross-validation within the training data to compute an estimate of the behavior decoding and neural self-prediction of each architecture using the training data. We then select one final architecture purely based on training data to be used for that cross-validation fold based on one of two criteria: 1) decoding focused: pick the architecture with the best neural self-prediction in training data among all those that reach within 1 s.e.m. of the best behavior decoding. 2) self-prediction focused: pick the architecture with the best behavior decoding in training data among all those that reach within 1 s.e.m. of the best neural self-prediction. The first criteria prioritizes good behavior decoding in the selection and the second criteria prioritizes good self-prediction. Note that these two criteria that are used when selecting among different learned models with different nonlinearities are completely independent of the internal objective functions used in learning the parameters for a given model (**S Fig. 1**). For example, in stage one of RNN PSID, RNN model parameters for a given model are always optimized to achieve good behavior decoding. But when selecting among different learned models with different combinations of nonlinearities, if desired one could perform that selection based on neural self-prediction. Thus, whenever neural self-prediction is also of interest, we report the results for both criteria (e.g. **Figs. 3 and 4** and **S Figs. 3 and 4**).

When comparing models with nonlinearity in different individual parameters to find the parameter that is largely sufficient for compressing nonlinearities (**Fig. 4**), we only consider one network architecture for the nonlinearity and that is having one hidden layer with 64 units.

### Numerical simulations

To validate RNN PSID in numerical simulations, we perform two sets of simulations one validating linear modeling and to show the correctness of the two-stage learning approach with numerical optimization and one validating nonlinear modeling. In the linear simulation, we randomly generate 100 linear models with various dimensionality and noise statistics, as described in our prior work^1^. Briefly, the neural and behavior dimensions are selected from 5 ≤ *n*_*y*_, *n*_*z*_ ≤ 10, the state dimension is selected from 1 ≤ *n*_*x*_ ≤ 10, and the number of latent state dimensions driving behavior is selected from 1 ≤ *n*_1_ ≤ *n*_*x*_, with all selections being random with uniform probability. Eigenvalues of the state transition matrix are selected randomly as complex conjugate pairs with uniform probability within the unit disk. Each element in the behavior and neural readout matrices is generated as a random Gaussian variable. State and neural observation noise covariances are generated as random positive definite matrices, and then scaled randomly with a number between 0.01 to 100 to obtain a wide range of relative noises across random models. A separate random linear state-space model with up to 10 latent state dimensions is generated to produce the behavior readout noise *∈*_*k*_, representing the behavior dynamics that are not encoded in the recorded neural activity. Finally, the behavior readout matrix is scaled to set the ratio of the signal s.d. to noise s.d. in each behavior dimension to a random number from 1 to 100. We perform model learning and evaluation with 2-fold cross-validation (**S Fig. 2**).

In the nonlinear simulations that are used to validate both RNN PSID and the hypothesis testing procedure it enables to find the origin of nonlinearity, we generate 20 random linear scalar models with *n*_*y*_ = *n*_*z*_ = *n*_*x*_, and then replace one of the four model parameters (i.e. *A*′, *K*, *C*_*y*_, and *C*_*z*_) with a nonlinear trigonometric function, such that roughly one period of the trigonometric function is visited by the model. To do this, we first scale the latent state in the initial random scaler linear model to find a similarity transform for it where the latent state has a 95% confidence interval range of 2π. We then add a sine function to the original parameter that is to be changed to nonlinear and scale the amplitude of the sine such that the amplitude of sine function reaches roughly 0.25 of the range of the outputs from the original linear parameter. This was done to reduce the chance of generating unrealistic unstable nonlinear models that produce outputs with infinite energy, which is likely when *A*′ is nonlinear. Changing one parameter to nonlinear can change the range of the statistics of the latent states in the model, thus we generate some simulated data from the model and redo the scaling of the nonlinearity until ratio conditions are met.

To generate data from the nonlinear model in equation (1), we first generate a neural noise time series *e*_*k*_ based on its covariance *Σ*_*e*_ in the model and initialize the state as *x*_0_ = 0. We then iteratively apply the second and first lines of equation (1) to get the simulated neural activity *y*_*k*_ from line 2 and then the next state *x*_*k*+1_ from line 1, respectively. Finally, once the state time series is produced, we generate a behavior noise time series *∈*_*k*_ based on its covariance *Σ*_*∈*_ in the model, and apply the third line of equation (1) to get the simulated behavior *z*_*k*_. Similar to linear simulations, we perform the modeling and evaluation of nonlinear simulations with 2-fold cross-validation (**S Fig. 3**).

### Neural datasets and behavioral tasks

We investigate the nonlinearities in four datasets with different behavioral tasks, brain regions, and neural recording modalities to show the generality of our methods and conclusions. Across all datasets, the spiking activity was binned with 10 ms nonoverlapping bins, smoothed with a Gaussian kernel with a 50 ms s.d., and then downsampled to 50 ms to be used as the neural signal to be modeled. The behavior time series was also downsampled to a matching 50 ms before modeling. In the three datasets where LFP activity was also available, we also studied two types of features extracted from LFP. As the first LFP feature, we considered raw LFP activity itself, which was low-pass filtered below 10 Hz (i.e. anti-aliasing) and downsampled to the behavior sampling rate of 50 ms timestep (i.e. 20 Hz). As the second feature, we computed the LFP log-powers in 8 standard frequency bands (delta: 0.1-4 Hz, theta: 4-8 Hz, alpha: 8-12 Hz, low beta: 12-24 Hz, high beta: 24-34 Hz, low gamma: 65-95 Hz, and high gamma: 130-170 Hz), in sliding 300 ms windows at a time step of 50 ms using Welch’s method (using 8 subwindows with 50% overlap)^1^.

### First dataset: 3D reaches to random targets

In the first dataset, the monkey (monkey J) performed reaches to a target randomly positioned in 3D space within the reach of the monkey, grasped the target, and then retuned its hand to resting position^1,26^. Angles of 27 joints in the shoulder, elbow, wrist, and fingers in the active hand (right hand) were tracked using 3D markers and taken as the behavior signal^1,26^. Neural activity was recorded with a 137-electrode microdrive (Gray Matter Research), out of which 28 electrodes were in the contralateral primary motor cortex M1. The multiunit spiking activity in these M1 electrodes was used as the neural signal. For LFP analyses, LFP features were also extracted from the same M1 electrodes. We analyzed the data from 7 recording sessions.

To visualize the low-dimensional latent state trajectories for each behavioral condition (**Fig. 6**), we determined the periods of reach and return movements in the data, resampled them to have similar number of time samples and averaged the latent states across those resampled trials. Given the redundancy in latent descriptions (i.e. any scaling, rotation, etc. on the latent states still gives an equivalent model), before averaging trials across cross-validation folds and sessions we devised the following procedure to standardize the latent states for each fold in the case of 2D latent states (**Fig. 6**): 1) We z-score all state dimensions to have zero mean and unit variance. 2) We rotate the 2D latent states such that the average 2D state trajectory for the first condition (here the reach epochs) starts from an angle of 0. 3) We estimate the direction of the rotation for the average 2D state trajectory of the first condition, and if it is not counterclockwise, we multiply the second state dimension by −1 to make it so. Note that in each step, the same mapping is applied to the latent states during the whole test data, regardless of condition, so this procedure does not alter the relative differences in the state trajectory across different conditions. The procedure also does not change the learned model and simply corresponds to a similarity transform that changes the basis of the model. All in all, this procedure only removes the redundancies for describing a 2D latent state-space model and standardizes the extracted latent states so that trials across different test sets can be averaged together.

### Second dataset: saccadic eye movements

In the second dataset, the monkey (monkey A) performed saccadic eye movements to one of eight targets on a display^28,29^. The 2D position of the eye was tracked and taken as the behavior signal. Neural activity was recorded with a 32-electrode microdrive (Gray Matter Research) covering the prefrontal cortex^1,27^. The single unit activity from these electrodes, ranging from 34 to 43 units across different recording sessions was used as the neural signal. For LFP analyses, LFP features were also extracted from the same 32 electrodes. We analyzed the data from the first 7 days of recordings. We only included data from successful trials where the monkey performed the task correctly by making a saccadic eye movement to the specified target. To visualize the low-dimensional latent state trajectories for each behavioral condition (**Fig. 6**), we grouped the trials based on their target position. Standardization across folds before averaging was done as in the first dataset.

### Third dataset: sequential reaches with a 2D cursor controlled with a manipulandum

In the third dataset, which was collected and made publicly available by the Miller lab^28,29^, the monkey (monkey T) controlled a cursor on a 2D screen using a manipulandum and performed a sequential reach task^28,29^. The 2D cursor position and velocity were taken as the behavior signal. Neural activity was recorded using a 100-electrode microelectrode array (Blackrock Microsystems, Salt Lake City, UT) in the dorsal premotor cortex (PMd)^28,29^. The single unit activity, recorded from 37 to 49 units across recording sessions, was used as the neural signal. This dataset did not include any LFP recordings, so LFP features could not be considered. We analyzed the data from all 3 recording sessions. To visualize the low-dimensional latent state trajectories for each behavioral condition (**Fig. 6**), we grouped trials into 8 different conditions based on the angle of the direction of movement (i.e. end position minus starting position) during the trial, with each condition covering movement directions within a 45 (i.e. 360/8) degree range. Standardization across folds before averaging was done as in the first dataset.

### Fourth dataset: random reaches with a 2D cursor displayed in virtual reality and controlled with the fingertip

In the fourth dataset, which was collected and made publicly available by the Sabes lab^30^, the monkey (monkey I) controlled a cursor on a 2D surface within a 3D virtual reality environment^18,30^. The cursor was controlled based on the fingertip position of the monkey^18,30^. The 2D cursor position and velocity were taken as the behavior signal. Neural activity was recorded with a 96-electorde microelectrode array (Blackrock Microsystems, Salt Lake City, UT)^18,30^ covering M1. We selected a random subset of 32 of these electrodes, which had 77 to 99 single units across the recording sessions, as the neural signal. For LFP analyses, LFP features were also extracted from the same 32 electrodes. We analyzed the data for the first seven sessions for which the wideband activity was also available (sessions 20160622/01 to 20160921/01). Grouping into conditions for visualization of low-dimensional latent state trajectories (**Fig. 6**) was done as in the third dataset. Standardization across folds before averaging was done as in the first dataset.

### Statistics

We used the Wilcoxon signed-rank test for all paired statistical tests.

## Data availability

Two of the datasets used in this work are publicly available^28–30^. The other datasets used to support the results are available upon reasonable request from the corresponding author.

## Code availability

The code for RNN PSID will be online at https://github.com/ShanechiLab/PyPSID.

## Acknowledgements

This work was supported in part by the following organizations and grants: the Office of Naval Research (ONR) Young Investigator Program (YIP) under contract N00014-19-1-2128 and NIH Director’s New Innovator Award DP2-MH126378 (to M.M.S.) and a University of Southern California Annenberg Fellowship (to O.G.S).

## Author contributions

O.G.S. and M.M.S. conceived the study, developed the new PSID algorithm, and wrote the manuscript, and O.G.S. performed all the analyses. B.P. provided two of the nonhuman primate datasets. M.M.S. supervised the work.

## Supplementary Figures

**S Fig. 1.**
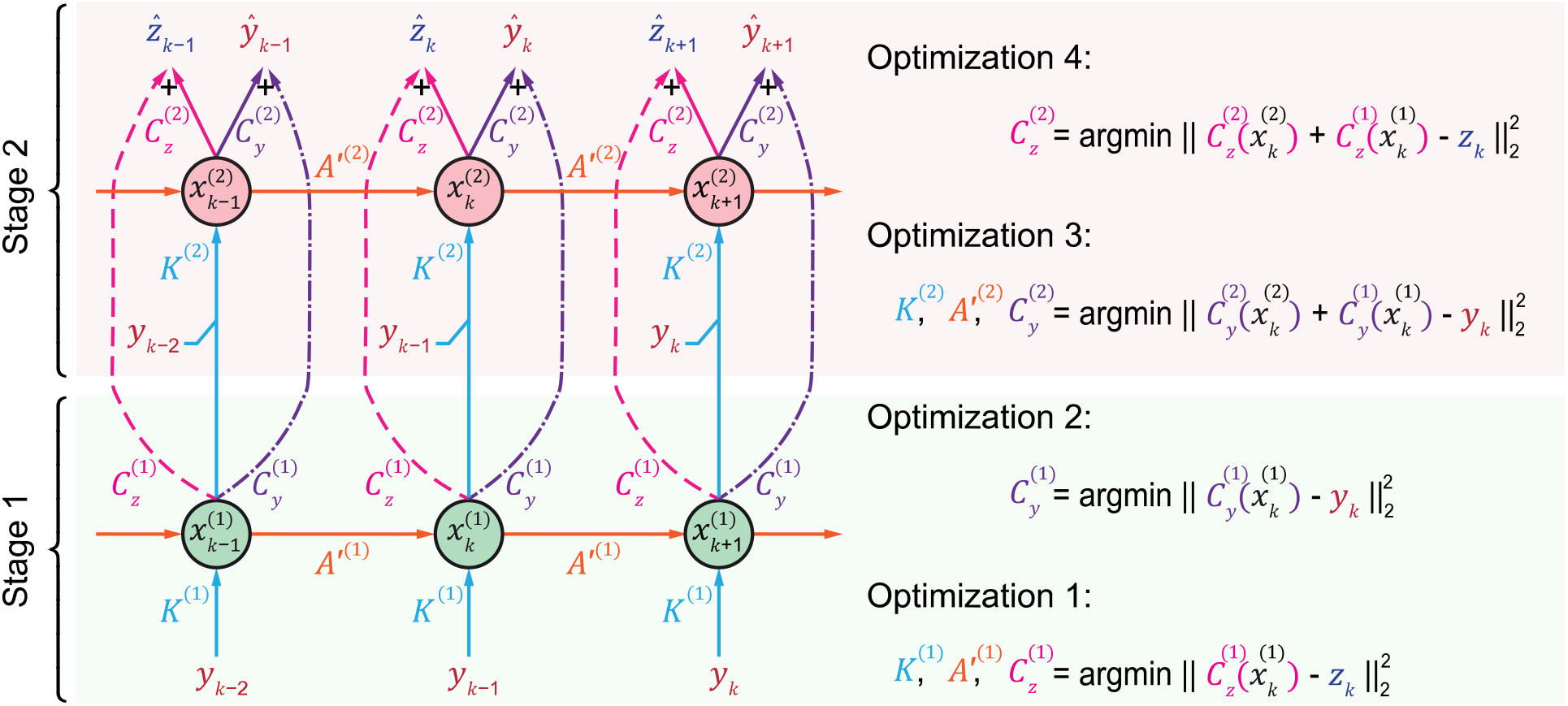
Detailed computation graph for both stages of RNN PSID. When both stages of RNN PSID are used, the computation graph is as shown in the figure. The learning consists of 4 numerical optimization problems (Methods): 1) Learn *A*′^(1)^, *K*^(1)^, and 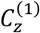 by fitting an RNN that minimizes behavior prediction error; 2) Learn 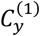 by fitting a feed-forward neural network that minimizes neural prediction error when using the RNN states 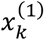 as input; 3) Learn *A*′^(2)^, *K*^(2)^, and 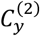 by fitting an RNN that minimizes the error in neural prediction when using the past neural activity and the states extracted from the first RNN 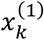 as input (note that 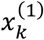 is computed in stage 1 and has known values for stage 2); and 4) Learn 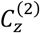 by fitting a feed-forward neural network that minimizes neural prediction error when using the second RNN states 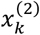 as input. Stage 1 consists of the first two and stage 2 consists of the latter two optimization problems. The dimension of 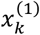 and 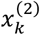 are hyperparameters that need to be determined by the user. The architecture of the feed-forward neural network that constructs each model parameter (**Fig. 1d-e**) can either be determined exactly by the user (e.g. for linear PSID and for individual nonlinearities in **Fig. 4**) or can be automatically selected among a range of architectures using an inner-cross-validation within the training data (e.g. flexible nonlinearity in **Figs. 3–4**, Methods).

**S Fig. 2.**
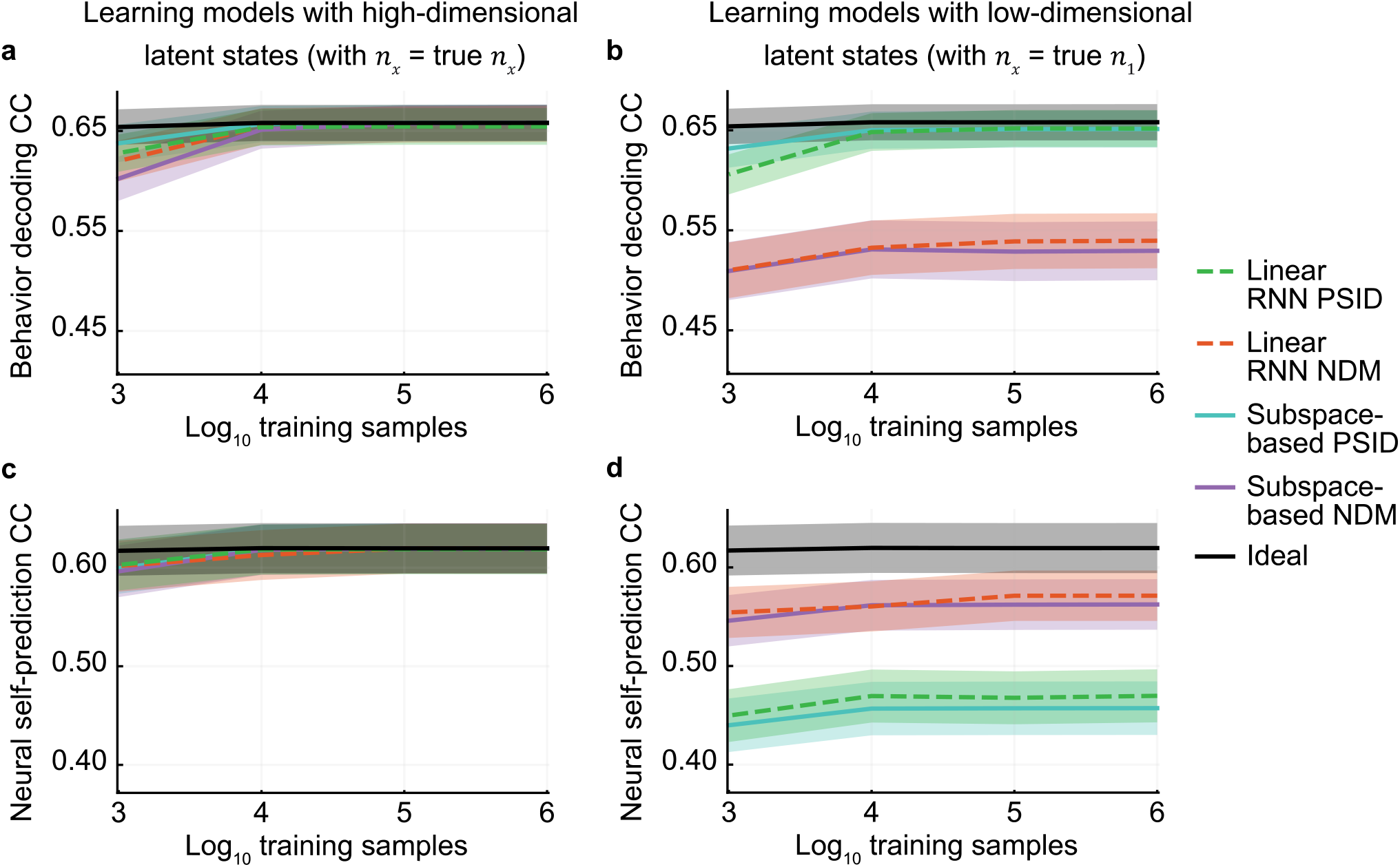
RNN PSID dissociates and prioritizes the behaviorally relevant neural dynamics while also learning the other neural dynamics in numerical simulations of linear models. (**a**) Cross-validated behavior decoding accuracy (correlation coefficient, CC) for each method as a function of the number of training samples when we use a state dimension equal to the total state dimension in the true model. Performance for the true model is shown in black. Solid lines show the mean across 100 random models and the shaded area shows the s.e.m. (**b**) Same as (a), but when learned models have low-dimensional latent states with enough dimensions just for the behaviorally relevant latent states (i.e. *n*_*x*_= *n*_1_). (**c-d**) Same as (a-b), showing the cross-validated neural self-prediction accuracy. Linear RNN PSID, just like subspace-based PSID^1^, achieves almost ideal behavior decoding even with low-dimensional latent states (panel b) showing that it correctly dissociates and prioritizes behaviorally relevant dynamics, while still being able to learn overall neural dynamics accurately if state dimension is high enough (panel c).

**S Fig. 3.**
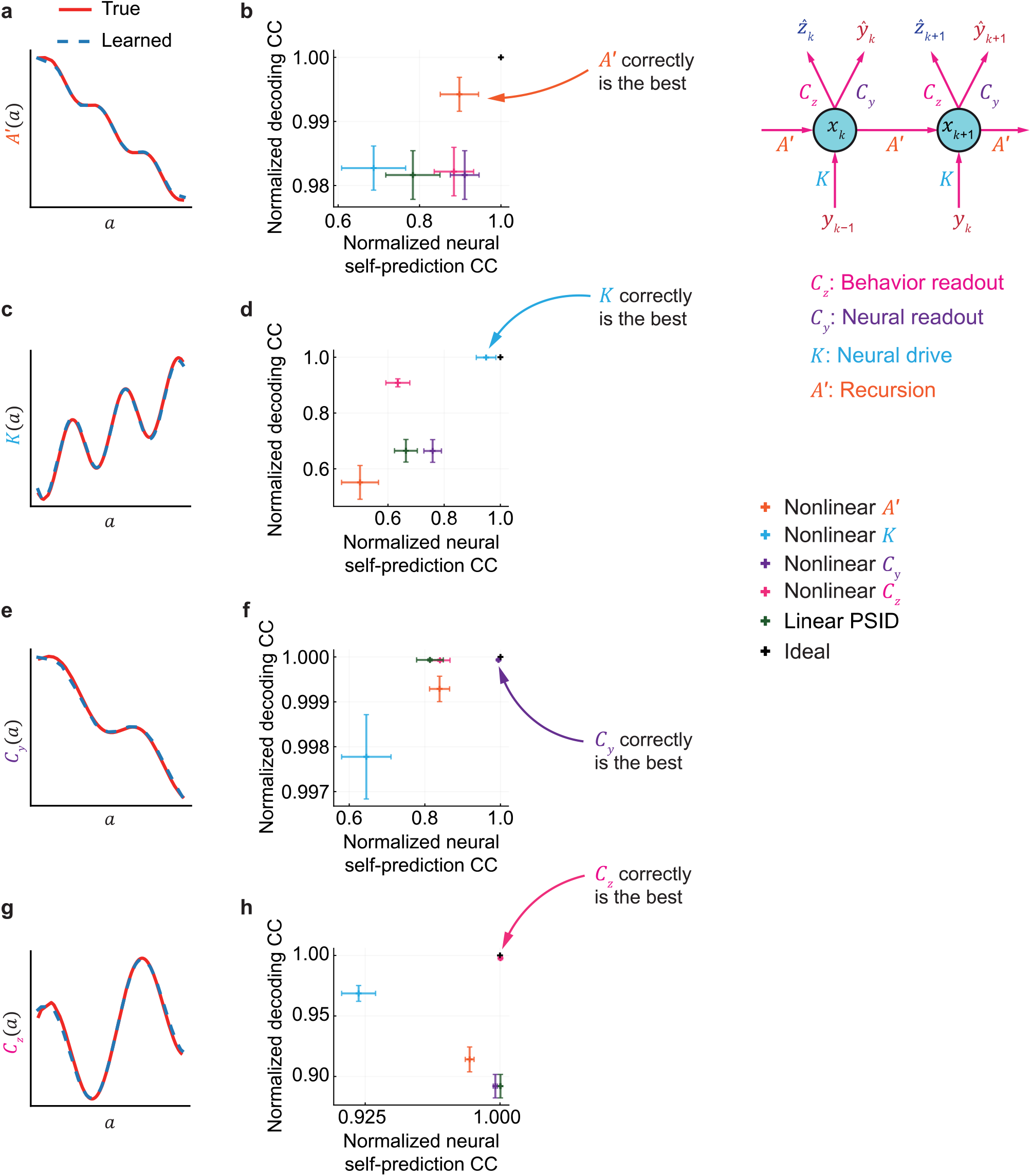
RNN PSID successfully identifies the origin of nonlinearity and learns it in numerical simulations. (**a**) Example true value for nonlinear recursion parameter *A*′ and the nonlinear value that RNN PSID learned for a random model for which only *A*′ was nonlinear. (**b**) Behavior decoding and neural self-prediction accuracy achieved by each type of nonlinearity when the true random models only had nonlinearity is in the recursion parameter *A*′. The performance measures for each random model are normalized by their ideal values that were achieved by the true model itself. Horizontal and vertical whiskers show the s.e.m. for neural self-prediction and behavior decoding, respectively. The model whose (neural self-prediction, behavior decoding) performance pair is at the upper-right corner of the plots has both the best behavior decoding and the best neural self-prediction and is thus chosen as the model that specifies the origin of nonlinearity. Among all types of nonlinearities, the correct one (i.e. *A*′) achieves the best performance overall, suggesting that by fitting and comparing RNN PSID models with different nonlinearities we can correctly find the origin of nonlinearity in the data. (**c-d**) Same as (a-b), for models that only have nonlinearity in the neural drive parameter *K*. (**e-f**) Same as (a-b), for models that only have nonlinearity in the behavior readout parameter *C*_*y*_. (**g-h**) Same as (a-b), for models that only have nonlinearity in the neural readout parameter *C*_*z*_.

**S Fig. 4.**
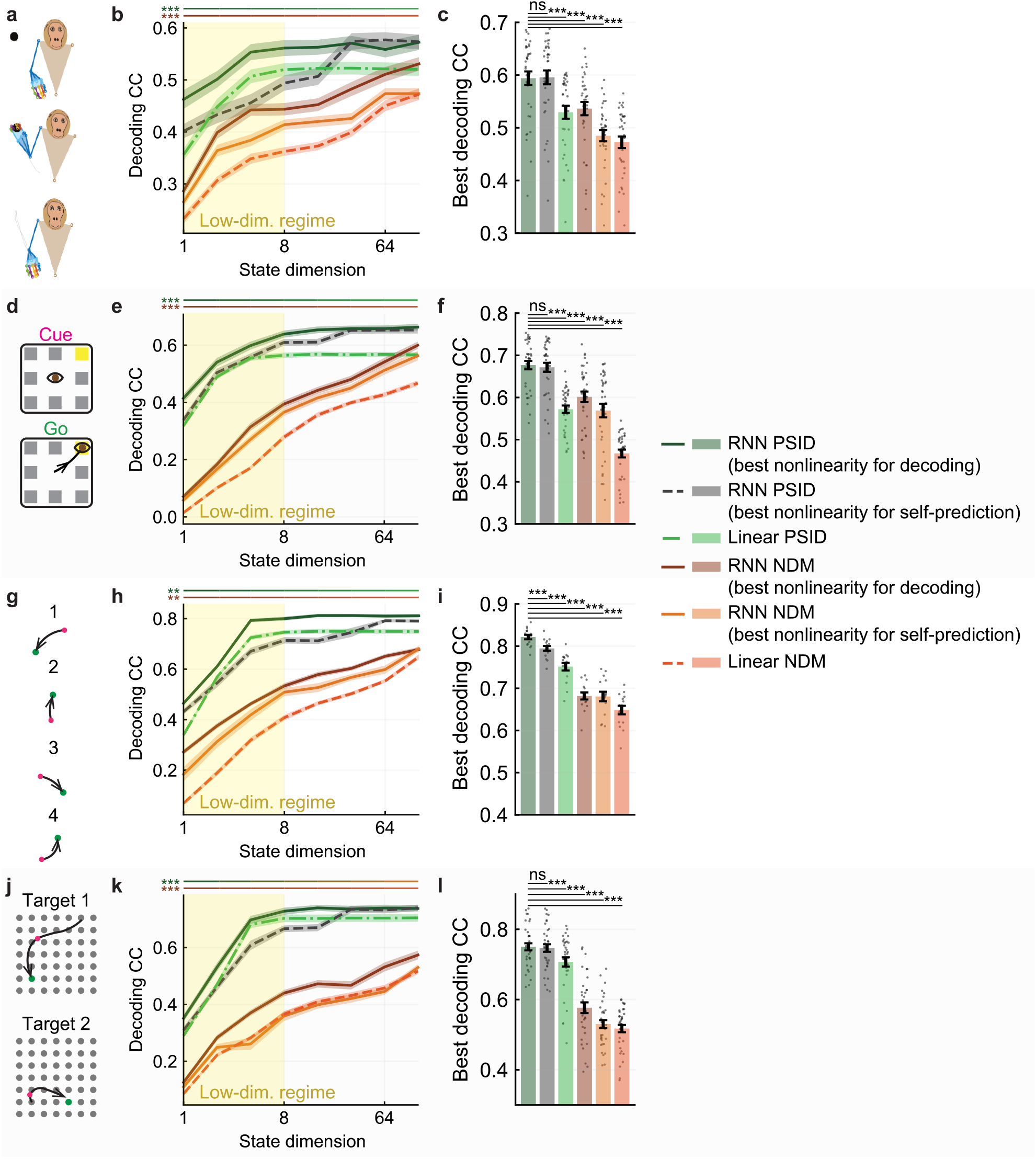
Behavior decoding of RNN PSID when using both stages shows that nonlinear RNN PSID outperforms linear PSID and other methods across all latent state dimensions, including high dimensions. (**a**) The 3D reach task. (**b**) Cross-validated behavior decoding accuracy (CC) achieved by variations of nonlinear and linear RNN PSID/NDM, for different latent state dimensions. Notation is as in **Fig. 2b**. Across latent state dimensions, the statistical significance of a one-sided pairwise comparison between nonlinear PSID/NDM (with best nonlinearity for decoding) vs linear PSID/NDM is shown with a horizontal green/red line with asterisks next to it (*N* = 35). For all latent state dimensions, nonlinear modeling results in significantly more accurate behavior decoding. (**c**) Peak behavior decoding accuracy achieved by each method, by choosing the state dimension in each session and fold as the smallest that reaches peak decoding accuracy. Bars, whiskers, and dots are defined as in **Fig. 2c**. (**d**-**f**) Same as (a-c) for the second dataset, with saccadic eye movements (*N* = 35). (**g**-**i**) Same as (a-c) for the third dataset, with sequential cursor reaches controlled via a 2D manipulandum (*N* = 15). (**j**-**l**) Same as (a-c) for the fourth dataset, with random grid virtual reality cursor reaches controlled via fingertip position (*N* = 35). For all RNN PSID variations, the first 16 latent state dimensions are learned using stage 1 and the remaining are learned using stage 2 (i.e. n_1_ = 16).

**S Fig. 5.**
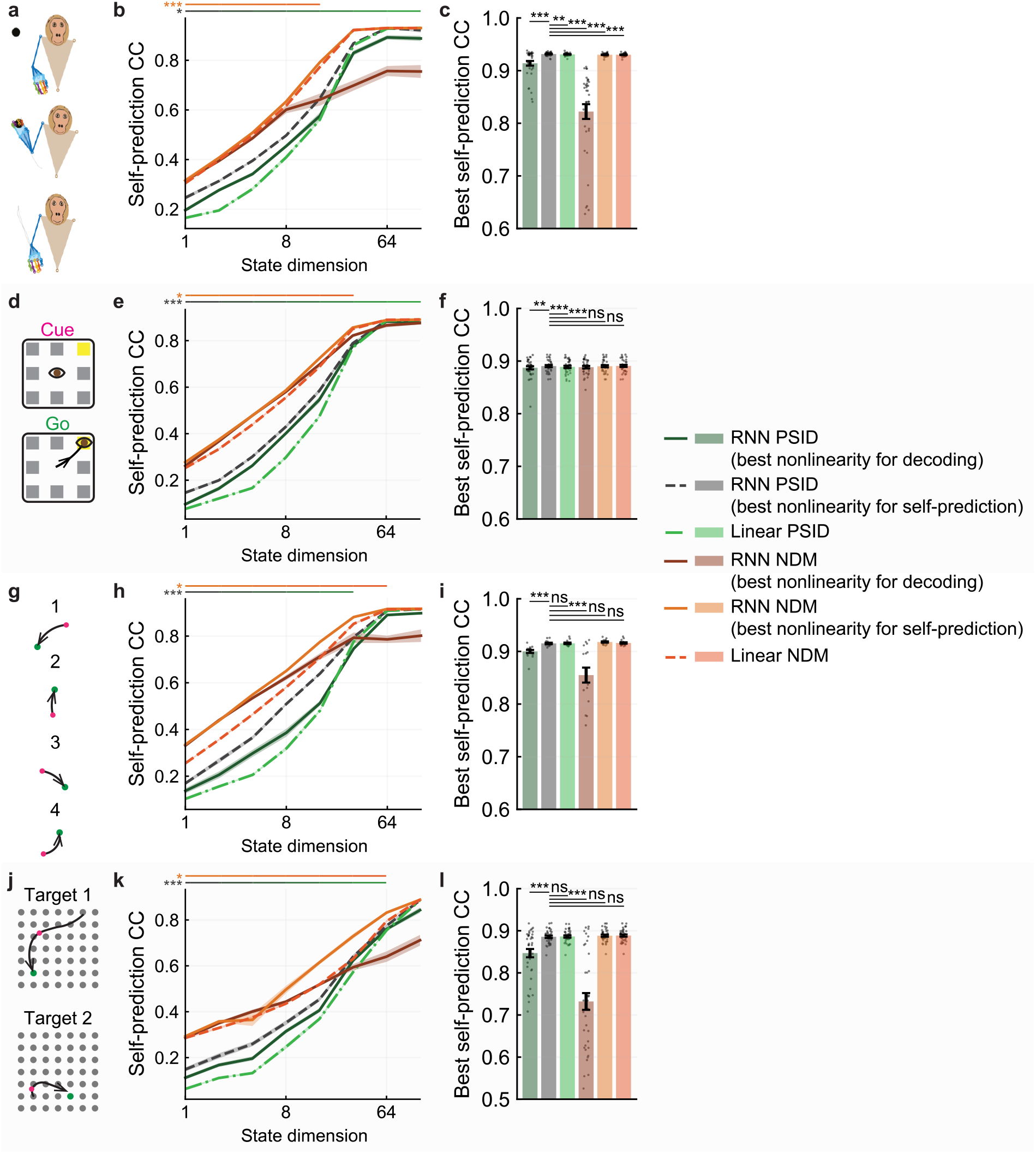
Neural self-prediction of RNN PSID when using both stages demonstrates that linear models can reach similar neural self-prediction accuracy as nonlinear models but only with high enough latent state dimensions. (**a**) The 3D reach task. (**b**) Cross-validated neural self-prediction accuracy (CC) achieved by variations of nonlinear and linear RNN PSID/NDM, for different latent state dimensions. Notation is as in **Fig. 2b**. Across latent state dimensions, the statistical significance of a one-sided pairwise comparison between nonlinear PSID/NDM (with best nonlinearity for self-prediction) vs linear PSID/NDM is shown with a horizontal black/orange line with asterisks next to it (*N* = 35). For low dimensional latent states, nonlinear modeling results in significantly more accurate neural self-prediction. (**c**) Peak neural self-prediction accuracy achieved by each method when choosing the state dimension in each session and fold as the smallest that reaches peak neural self-prediction. Bars, whiskers, and dots are defined as in **Fig. 2c**. (**d**-**f**) Same as (a-c) for the second dataset, with saccadic eye movements (*N* = 35). (**g**-**i**) Same as (a-c) for the third dataset, with sequential cursor reaches controlled via a 2D manipulandum (*N* = 15). (**j**-**l**) Same as (a-c) for the fourth dataset, with random grid virtual reality cursor reaches controlled via fingertip position (*N* = 35). For all RNN PSID variations the first 16 latent state dimensions are learned using stage 1 and the remaining are learned using stage 2 (i.e. n_1_ = 16).

**S Fig. 6.**
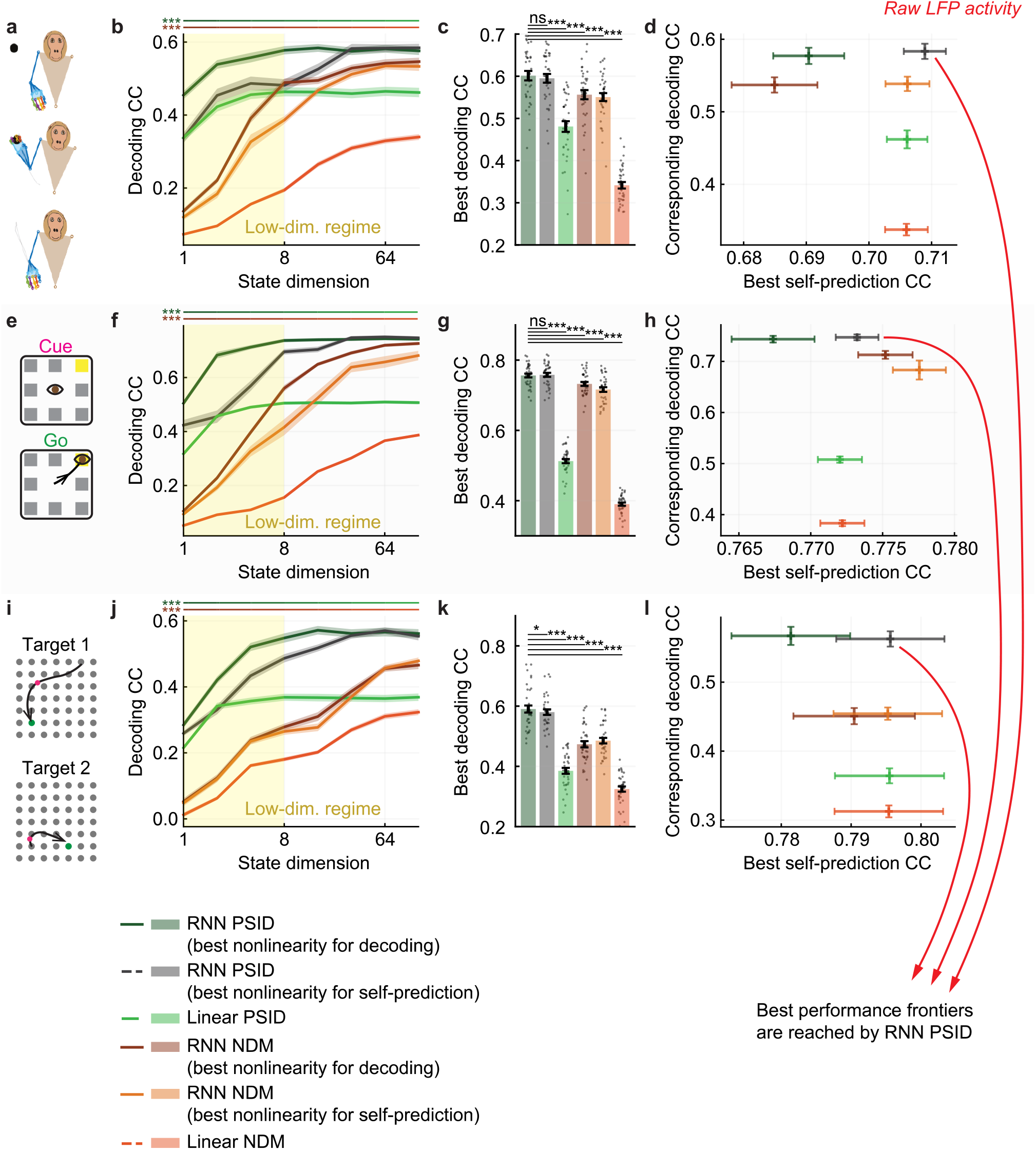
For raw LFP activity, RNN PSID more accurately learns behaviorally relevant neural dynamics while also reaching similar neural self-prediction accuracy as RNN NDM. Figure content is parallel to **S Fig. 4** and **Fig. 3**, shown for raw LFP activity. (**a**) The 3D reach task. (**b**) Cross-validated behavior decoding accuracy (CC) achieved by variations of nonlinear and linear RNN PSID/NDM, for different latent state dimensions. Notation is as in **S Fig. 4b**. (**c**) Peak behavior decoding accuracy achieved by each method, by choosing the state dimension in each session and fold as the smallest that reaches peak decoding accuracy. Bars, whiskers, and dots are defined as in **Fig. 2c**. (**d**) Peak neural self-prediction accuracy achieved by each method shown on the horizontal axis versus the corresponding behavior decoding accuracy on the vertical axis. Latent state dimension for each method in each session and fold is selected as the smallest that reaches peak neural self-prediction, thus peak decoding accuracies are not exactly the same as in (c). Notation is as in **Fig. 3**, with the plus on the plot showing the mean self-prediction and decoding accuracy across sessions and folds (*N* = 35), and the horizontal and vertical whiskers showing the s.e.m. (**e**-**h**) Same as (a-d) for the second dataset, with saccadic eye movements (*N* = 35). (**i**-**l**) Same as (a-d) for the fourth dataset, with random grid virtual reality cursor reaches controlled via fingertip position (*N* = 35).

**S Fig. 7.**
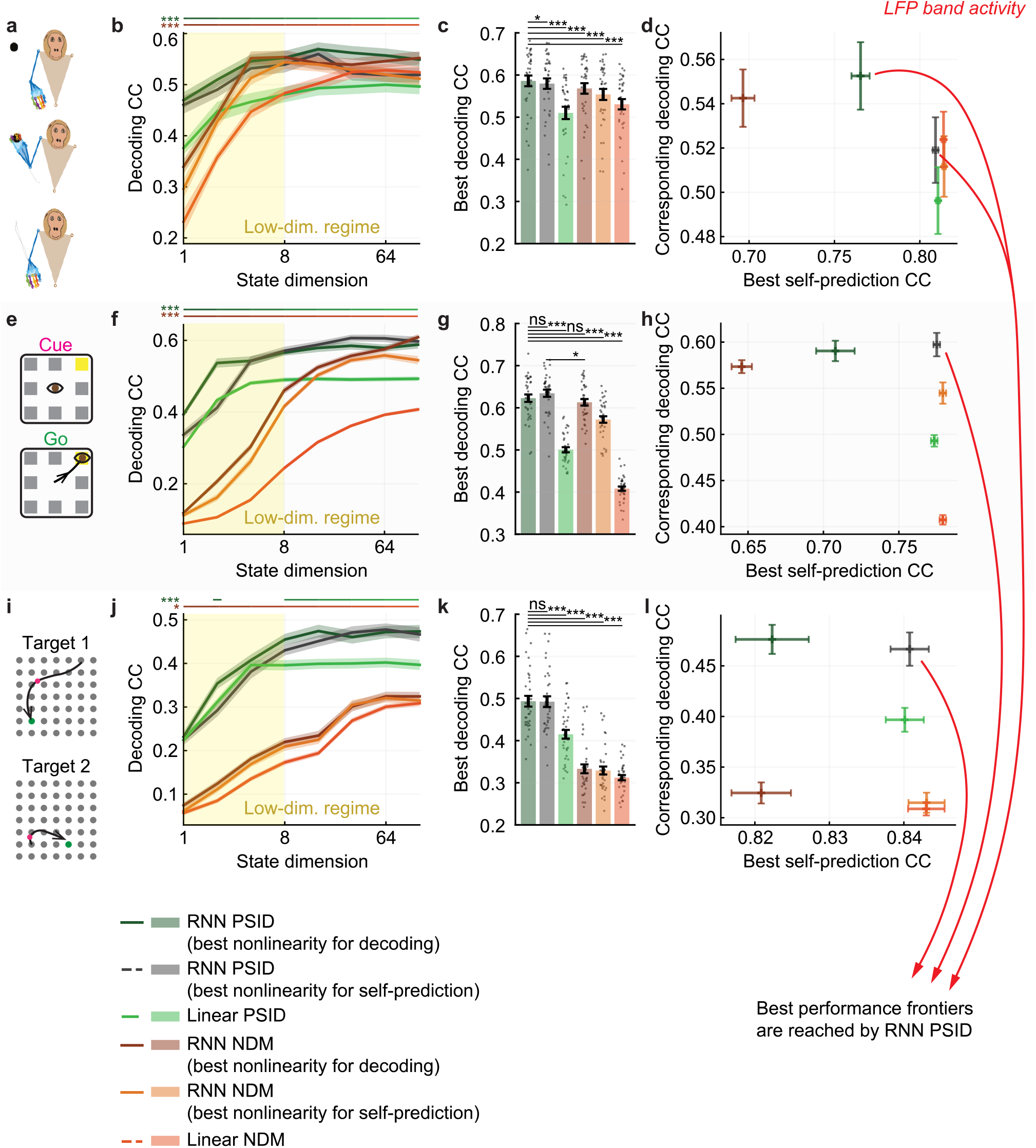
For LFP band power activity, RNN PSID more accurately learns behaviorally relevant neural dynamics while also reaching similar neural self-prediction accuracy as RNN NDM. Figure content is exactly parallel to **S Fig. 6**, shown for LFP band power activity.

**S Fig. 8.**
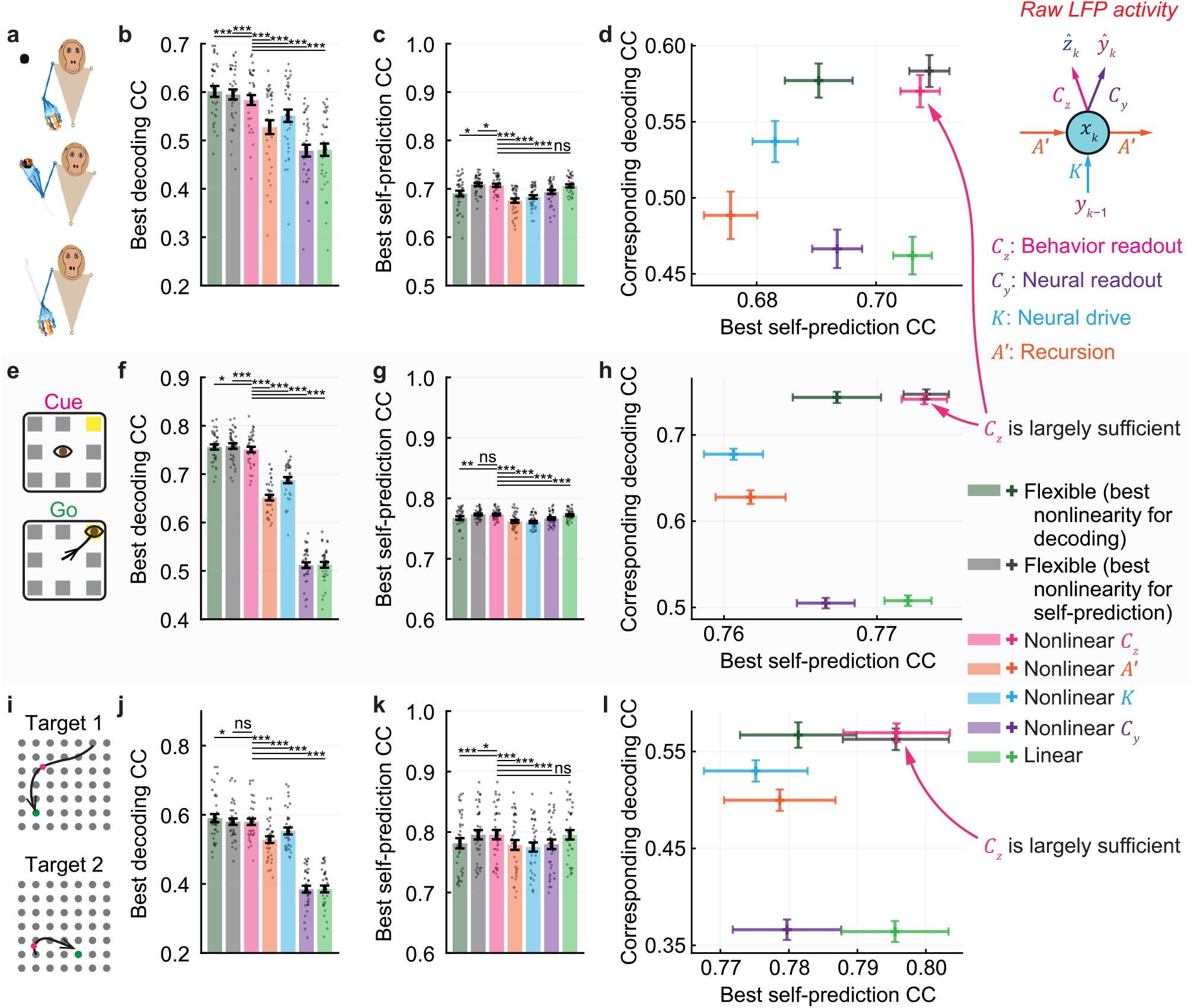
For raw LFP activity, RNN PSID again reveals that nonlinearities can be largely captured by the behavior readout of the model across datasets. Figure content is exactly parallel to **Fig. 4**, shown for raw LFP activity. The behavior readout nonlinearity does better than every other individual nonlinearity and comparable to when nonlinearity is flexibly chosen to be in all or any combination of parameters.

**S Fig. 9.**
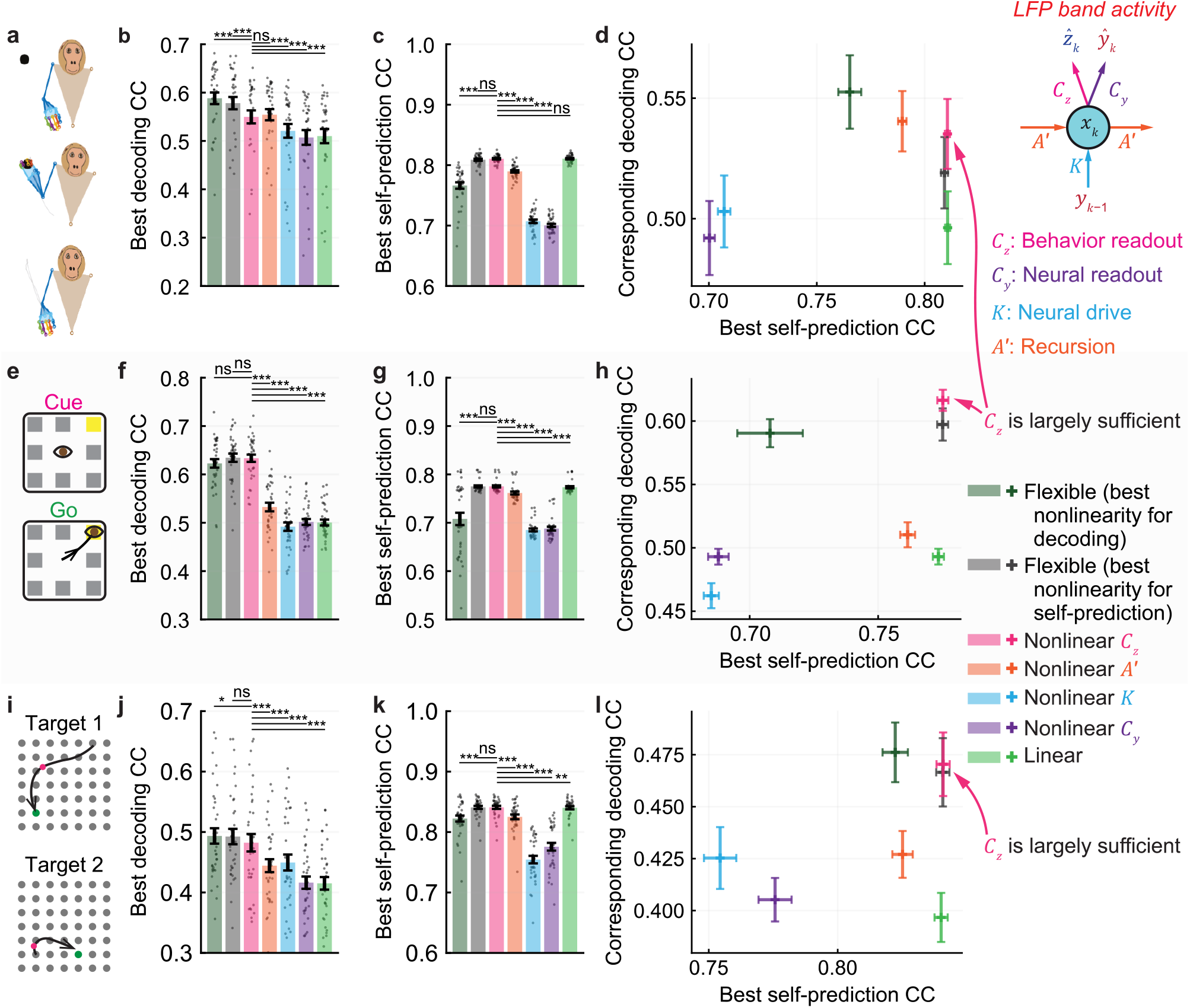
For LFP band power activity, RNN PSID again reveals that nonlinearities can be largely captured by the behavior readout of the model across datasets. Figure content is exactly parallel to **Fig. 4**, shown for LFP band power activity. The behavior readout nonlinearity does better than every other individual nonlinearity and comparable to when nonlinearity is flexibly chosen to be in all or any combination of parameters.

## Supplementary Notes

### S Note 1: Relation of RNN PSID to linear state-space models in predictor form

The model used by RNN PSID is provided in equation (1). Here, we expand on the motivation behind this model formulation and its relation to linear state-space models. As a general linear model, neural activity 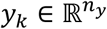 and behavior 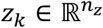 can be jointly modeled as

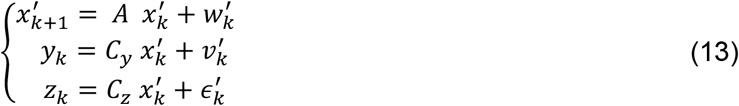

where 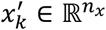 is a latent variable, 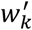 and 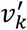 are Gaussian white noises, and 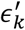 is a general random process that is independent of neural activity and represents any behavior dynamics that are not encoded in neural activity^1^. Given the above linear model, the latent states can be estimated from the neural activity *y*_*k*_ using a Kalman filter

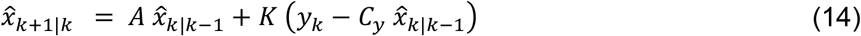

where *K* is the Kalman gain^21,39^. Equation (13) can be equivalently written in terms of the Kalman estimated states 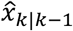 as

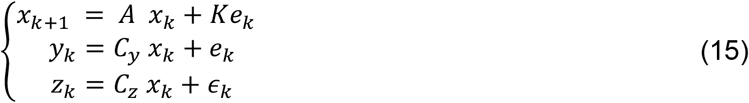

where 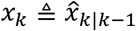 and *e*_*k*_ and *∈*_*k*_ are respectively the parts of neural and behavior signals that cannot be predicted from past neural activity (i.e. 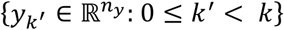). Equivalently, by replacing *e*_*k*_ from the first line with its value from the second line, we can also write equation (15) as

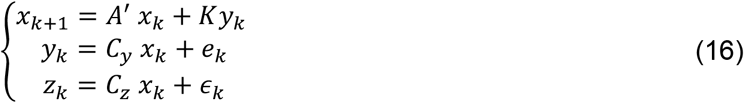

where 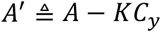. Equations (13) and (16) describe the same second order statistics for the observed times series *y*_*k*_ and *z*_*k*_ and thus are equivalent^21,39^. These two formulations are referred to as the stochastic and predictor forms, respectively^21,39^.

In equation (16), each multiplication between a model parameter and a vector (e.g. *A*′*x*_*k*_) can be thought of as a multi-input-multi-output linear function applied to an input vector (e.g. function *A*′(.), applied to *x*_*k*_). Rewriting all matrix multiplications as multi-input-multi-output functions we get

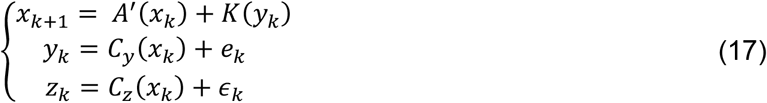

where each function (e.g. *A*′(.)) is a parameter to be learned. This gives the model form for the RNN PSID model in equation (1). Unlike the linear state-space model, however, RNN PSID has the additional generalization that we can allow any subset of the parameters in its model to be general nonlinear functions.

Given that the model in equation (1) is constructed in the predictor form, even when parameters are nonlinear, the model can still be readily used to estimate the latent states *x*_*k*_ given the neural observations *y*_*k*_, and to decode behavior *z*_*k*_. To do this, we run the first line of equation (1) to estimate the latent state and then pass this state through the learned *C*_*y*_ or *C*_*z*_ functions in the second or the third line to predict neural activity or decode behavior, respectively.

